# Untargeted metabolomics reveals key metabolites and genes underlying salinity tolerance mechanisms in maize

**DOI:** 10.1101/2025.04.08.647850

**Authors:** Manwinder S. Brar, Amancio De Souza, Avineet Ghai, Jorge F.S. Ferreira, Devinder Sandhu, Rajandeep S. Sekhon

## Abstract

Understanding the physiological, metabolic, and genetic mechanisms underlying salt tolerance is essential for improving crop resilience and productivity, yet their complex interactions remain poorly defined. We compared physiological and metabolic responses to salinity between two contrasting maize inbred lines: the salt-sensitive C68 and the salt-tolerant NC326. The senstitivity of C68 was characterized by reduced shoot and root dry weights and plant height, high tissue accumulation of Na and Cl, but low K, and lower leaf proline accumulation compared to the salt-tolerant NC326. Untargeted metabolomics identified 56 metabolites categorized as constitutively upregulated or salt-responsive. In NC326, constitutive accumulation of flavonoids, including schaftoside, tricin, and kaempferol-related compounds in leaves suggests adaptive priming against oxidative stress, while constitutively higher lipids and fatty acids in roots may enhance membrane stability. Salt-responsive metabolites, notably antioxidants and lanosterol, highlighted inducible oxidative-stress mitigation and membrane-stabilization strategies. By integrating metabolomic and genetic analyses, we identified 10 candidate genes involved in the biosynthesis of key metabolites. These findings establish a comprehensive platform for functional validation of metabolites and candidate genes for developing maize varieties with improved resilience to soil salinity through targeted breeding or biotechnological strategies.

**Plane Language Summary:** Salinity, or high salt content in soil, is a major challenge to crop growth worldwide, reducing food production. Maize, an important crop globally, struggles to grow under salty conditions. This study compared two maize types—one that grows well in salty soil and one that struggles—to understand how maize plants adapt to salt stress. Using advanced techniques, we measured hundreds of compounds (metabolites) in plant tissues to identify protective substances. We discovered that plants resistant to salt stress naturally produce higher amounts of protective substances, helping them avoid damage from salt. Additionally, we found specific metabolites that plants produce when exposed to salty conditions to protect their cells. We also identified genes that control the production of these important metabolites. This research provides new insights into how maize plants manage salt stress and highlights potential targets for developing crops that grow better in saline soils.

**Core Ideas:**

- Maize genotypes differ in growth, ion balance, and proline under salt stress.
- Metabolomics reveals pre-stress buildup of protective flavonoids and fatty acids.
- Salt triggers metabolic changes like sterol increase to protect membranes.
- Genetic analysis links genes to key metabolites controlling salt tolerance.
- Metabolite-gene insights guide breeding of maize for salt resilience

## Introduction

Salinity is a major abiotic stress that significantly reduces crop yields, affecting approximately 800 million hectares (6%) of global agricultural land and contributing to annual agricultural losses estimated between 12 and 27.3 billion USD (Butcher et al., 2016; Munns & Tester, 2008; Qadir et al., 2014; Zhou et al., 2022). The impact of salinity on crops, water, and agricultural soils is expected to worsen due to climate change, unsustainable irrigation practices, soil degradation, and the expansion of irrigated farming. Rising temperatures and increased drought conditions accelerate evaporation, resulting in higher salt concentrations in the soils, while sea-level rise causes saltwater intrusion into coastal aquifers used to irrigate farmland. Additionally, poor irrigation management, excessive fertilizer use, inadequate drainage, urbanization, and deforestation contribute to soil salinization by altering natural water flows and exposing underground salts, further reducing agricultural productivity (Tarolli et al., 2024). Saline soil typically has a higher accumulation of sodium, potassium, calcium, magnesium, and chloride ions. While salinization and sodification affect all climatic regions, arid and semi-arid climate zones are particularly affected due to limited natural leaching (Gheyi et al., 2023; Rengasamy, 2006). Salt stress disrupts essential physiological processes including photosynthesis, respiration, and water uptake, leading to significant losses in crop productivity (Ferreira et al., 2024). Given the increasing severity of soil salinity and its impact on agriculture, improving salt tolerance in crop plants is crucial for developing climate-resilient varieties and mitigating yield losses.

Maize (*Zea mays* L.), one of the most widely grown cereals in the world, is a multi-purpose crop that provides food, livestock feed, and various industrial products including biofuels, alcohol, and vegetable oil. Abiotic stresses including drought and salinity significantly reduce maize productivity and threaten global food security, feed availability, and industrial supply chains (Farooq et al., 2015; Nepolean et al., 2018; Sandhu et al., 2020). Maize is moderately sensitive to salinity with high soil NaCl concentrations leading to severe leaf wilting and stunted plant growth (Maas & Hoffman, 1977; Menezes-Benavente et al., 2004). High Na concentration interferes with potassium uptake thereby disrupting stomatal opening, increasing evapotranspiration, and causing leaf necrosis (Fortmeier & Schubert, 1995). Moreover, salt stress results in higher production of reactive oxygen species that cause oxidative damage to the cells (de Azevedo Neto et al., 2006). Understanding the genetic basis of salt tolerance is essential for sustainable maize productivity through various mechanisms including improved ion homeostasis, osmotic balance, and antioxidant defense. Breeding and genetic engineering approaches, including marker-assisted breeding and transgenic modifications, offer promising strategies for developing salt-tolerant maize genotypes.

Metabolites represent the functional readout of cellular biochemistry and, therefore, underlie external plant phenotypes. Plants exposed to salinity undergo dramatic metabolic shifts to maintain basic metabolism and adapt to the stressful environment (Arbona et al., 2013; Bundy et al., 2009). The accumulation of certain primary and secondary metabolites enhances salt tolerance by alleviating oxidative stress and regulating osmotic balance. Since the metabolome is closely linked to the phenotype, characterizing global metabolic changes is crucial for identifying novel metabolites and mechanisms underlying salt tolerance. However, there are critical knowledge gaps in the salinity-metabolome nexus, including the identity of metabolites associated with the salinity response and their genetic regulation. Recent metabolomic studies have provided insights into metabolic variation and the underlying molecular responses to salinity (Han et al., 2023; Liang et al., 2021; Tareq et al., 2024; Wang et al., 2023; Widodo et al., 2009). However, a comprehensive understanding of metabolic changes across different tissues during salt stress and their contributions to salt tolerance in maize remains largely unexplored.

Our recent study on the characterization of natural variation for salt tolerance in maize germplasm revealed tremendous variation and identified salt-tolerant and salt-sensitive inbred lines (Sandhu et al., 2020). The current study investigates the physiological, ionic, and metabolic responses of NC326 and C68 inbred lines to salt stress to uncover the mechanisms underlying salt tolerance. Through a time-course analysis of leaf and root metabolome, this study provides mechanistic insights into tissue-specific metabolic perturbations and identify putative pathways and genes underlying salt tolerance. These findings enhance our understanding of the biological processes governing salt tolerance and provide a platform for development of salt tolerance maize and related grasses.

## Material and Methods

### Plant materials and growth conditions

The maize inbred lines used in this study, C68 (salt-sensitive) and NC326 (salt-tolerant), were selected based on the phenotypic evaluation of a maize diversity panel for salinity response (Sandhu et al., 2020). These inbred lines were evaluated under controlled greenhouse conditions using lysimeters of 120 cm length, 60 cm in width, and 50 cm in depth at the US Salinity Laboratory, Riverside, CA. A randomized complete block design was employed, with twelve seeds sown per inbred line and subsequently thinned to nine plants each. Three biological replicates were used in the experiment. Initially, plants were cultivated in half-strength Hoagland’s solution with an electrical conductivity (EC_iw_) of 1.46 dS m^-1^ for 2 weeks (Table 1). Subsequently, the solution for the plants designated for the salinity treatment was gradually increased to 16 dS m^-1^ over three days to avoid osmotic shock (Table 1). The moment when the target salinity concentration (16 dS m^-1^) was reached was designated as 0h. The control plants continued to grow in half-strength Hoagland’s solution (EC_iw_ = 1.46 dS m⁻¹). Given that plants under salt stress first undergo osmotic stress before acclimating to saline conditions, tissue samples were collected at two additional time points (24h and 48h after reaching the final salinity level) to effectively capture physiological and metabolic adjustments during acclimation. These samples were immediately frozen in liquid nitrogen and stored at -80 °C for metabolic analysis. The remaining plants were grown under their respective treatments for an additional three weeks.

**Table 1.**
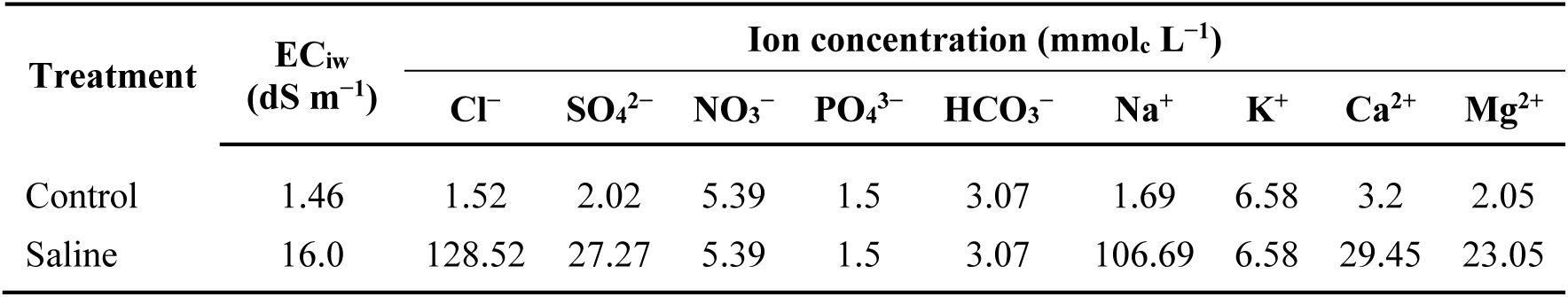
Composition of irrigation water.

| Treatment | EC<sub>iw</sub><br>(dS m<sup>-1</sup>) | Ion concentration (mmol<sub>c</sub> L<sup>-1</sup>) |  |  |  |  |  |  |  |  |
| --- | --- | --- | --- | --- | --- | --- | --- | --- | --- | --- |
|  |  | Cl<sup>-</sup> | SO<sub>4</sub><sup>2-</sup> | NO<sub>3</sub><sup>-</sup> | PO<sub>4</sub><sup>3-</sup> | HCO<sub>3</sub><sup>-</sup> | Na<sup>+</sup> | K<sup>+</sup> | Ca<sup>2+</sup> | Mg<sup>2+</sup> |
| Control | 1.46 | 1.52 | 2.02 | 5.39 | 1.5 | 3.07 | 1.69 | 6.58 | 3.2 | 2.05 |
| Saline | 16.0 | 128.52 | 27.27 | 5.39 | 1.5 | 3.07 | 106.69 | 6.58 | 29.45 | 23.05 |

### Data collection and sampling

Root and shoot tissues samples, obtained by separating plants at the scutellar node, were dried at 70 °C for 96 h to determine the dry root and shoot weight, respectively. Plant height was measured in centimeters (cm) from the tip of the primary root to the tip of the topmost visible leaf in the whorl.

### Mineral Analyses

Oven-dried leaf and root tissues were ground into a fine powder prior to mineral analyses. Chloride concentrations were determined using a colorimetric assay involving mercuric thiocyanate in the presence of ferric nitrate in an AQ300 discrete analyzer. Concentrations of other ions were measured after nitric acid digestion using inductively coupled plasma optical emission spectrometry (ICP-OES; 3300 DV, Perkin-Elmer Corp., Waltham, MA, USA).

### Proline Analysis

Proline extraction was carried out using 250 mg of finely ground, oven-dried leaf samples, each combined with 25 mL of deionized water. Samples were incubated in a water bath at 45 °C for one hour, with vigorous vortex mixing every 15 min (four times total). Following incubation, tubes were centrifuged at 5000 rpm, and the supernatant was filtered and used for proline determination. For each assay, 1 mL of leaf extract was combined with 1 mL acid ninhydrin reagent and 1 mL of glacial acetic acid. The mixture was shaken vigorously at 375 rpm for 12 min on a platform shaker (Innova 2000, New Brunswick Scientific, Edison, NJ), followed by heating in a boiling water bath (100 °C) for 1 hour to form the proline-ninhydrin chromophore complex. After cooling, the proline-ninhydrin complex was extracted by vigorous shaking with 2 mL toluene. The upper toluene layer containing the chromophore was transferred into a glass cuvette, and absorbance was measured at 520 nm using a spectrophotometer (DU 7500, Beckman Coulter, Brea, CA), with pure toluene serving as a blank. Proline concentrations were quantified using a standard curve and expressed as μmol g⁻¹ dry weight.

### Metabolite extraction

Freeze-dried leaf and root samples were ground to a fine powder using Geno/Grinder 2010 (SPEX SamplePrep, Metuchen, NJ, USA). For each sample, 10 mg of the powder was resuspended in 100 µL of a monophasic extraction solvent (30 acetonitrile:30 methanol:20 isopropanol:20 distilled water). The samples underwent sonication on ice for 5 min, followed by vortex-agitation for 90 min at 4°C, centrifugation at 4°C for 10 minutes at 21,255 G, and collection of supernatants into LC-MS vials. A pooled quality control (QC) sample, comprising equal volumes of all samples, was used to monitor system stability. LC-MS metabolomics analysis was performed at the UC Riverside Metabolomics Core Facility as described (Rothman et al., 2019), with minor modifications. Briefly, a Synapt G2-Si quadrupole time-of-flight mass spectrometer (Waters Corporation, Milford, MA, USA) coupled to an Acquity I-class UPLC system (Waters Corporation, Milford, MA, USA) was used, employing an Acquity CSH phenyl-hexyl column (2.1 mm x 100 mm, 1.7 µM) (Waters Corporation, Milford, MA, USA) for separations. The mobile phases consisted of (A) water with 0.1% formic acid and (B) acetonitrile with 0.1% formic acid. The column was maintained at 40°C with a flow rate of 250 µL/min and an injection volume of 1 µL. The gradient elution was as follows: 0 min, 1% B; 1 min, 1% B; 8 min, 40% B; 24 min - 26.5 min, 100% B; 27 min, 1% B; 30 min, 1% B. The MS data was acquired over a scan range of 50 to 1200 m/z with a 100 ms scan time, and MS/MS spectra were obtained in a data-dependent manner. The source temperature was set at 150°C, desolvation temperature at 600°C, with desolvation and cone gas at 600 L/hr and 0 L/hr, respectively. Nitrogen served as the desolvation gas, while argon was used as collision gas. The capillary voltage was maintained at 1 kV in positive-ion mode, and the leucine enkephalin pentapeptide (YGGFL) was infused for mass correction.

### Data processing

Untargeted LC-MS data processing, including peak picking, alignment, deconvolution, integration, normalization, and spectral matching was performed in Progenesis Qi software (Nonlinear Dynamics, Milford, MA, USA). Data was normalized using total ion abundance and the median, excluding features with variation greater than 30% across QC injections (Barupal et al., 2018; Dunn et al., 2011). To resolve multiple features annotated as the same metabolite, features were assigned to a cluster-ID using RAMClust (Broeckling et al., 2014). Annotation level confidence was assigned based on an extension of the Metabolomics Standard Initiative guidelines (Schymanski et al., 2014; Sumner et al., 2007), with level 1a indicating an MS, MS/MS, and retention time match to an in-house database, level 1b indicating an MS and MS/MS match to an in-house database, level 2a representing an MS and MS/MS match to an external database, level 2b denoting an MS and MS/MS match to the LipidBlast *in silico* database (Kind et al., 2013). Several mass spectral metabolite libraries were searched including Mass Bank of North America libraries, Metlin (Smith et al., 2005), and an in-house library, to enhance metabolite identification.

### Identification of genes regulating metabolites

Genes encoding enzymes catalyzing the biosynthesis of kaempferol and amyrin were obtained from CornCyc 10.0.1 (https://www.plantcyc.org) online database. Genes related to schaftoside, tricin, isovitexin, coumaroyl tyramine, and LPC 18:3 were found searching the online literature (Niu et al., 2023; Schmidt et al., 1999; Wang et al., 2020; Zhou et al., 2008). Gene for lanosterol biosynthesis was found by blasting the protein sequence of house mouse lanosterol synthase (BC029082.1) against maize protein database.

### Statistical Analysis

Statistical analysis was performed using a UCR Metabolomics Core proprietary tool, an HTML-based solution after metabolite identification and quantification. R packages prcomp was used to perform principal component analysis (PCA). Data was log-transformed for pairwise comparison between treatments, and differentially abundant metabolites were identified by applying Welch’s t-test. To control for multiple testing, P-values were adjusted using Benjamini and Hochberg correction to reduce false discovery rate.

## Results

### Distinct physiological responses of two inbred lines to salt stress

Exposure of two maize inbred lines to salt stress revealed that C68 (salt-sensitive) exhibited a significant decrease in shoot and root dry weight, whereas NC326 (salt-tolerant) had no significant change in these traits, as evident from comparison with the respective control plants and the salt tolerance index (Figure 1, A-B). Furthermore, while both inbred lines experienced a reduction in plant height under saline conditions, the decline was greater in C68 (68.5%) compared to NC326 (43.1%) (Figure 1C). These observations align with our previous findings that NC326 and C68 represent salt-tolerant and salt-sensitive inbred lines, respectively (Sandhu et al., 2020). To investigate if these phenotypic responses were associated with changes in ion concentrations, we analyzed sodium (Na), chloride (Cl), and potassium (K) concentrations in the leaves and roots of both inbred lines. Salt stress led to a significant increase in Na and Cl concentrations and a significant decrease in K concentration in both tissues (Figure 1, D-I). Notably, the relative increase in Na concentration was substantially higher in C68 across both tissues, whereas the relative increase in Cl concentration was greater in the leaves but comparable in roots between C68 and NC326. These results indicate that salinity perturbs the ion homeostasis to a greater extent in the C68 inbred line. Consistent with the positive association of proline accumulation and salt tolerance (El Moukhtari et al., 2020), NC326 accumulated significantly more leaf proline than C68 under salt stress, indicating that genetic differences in the synthesis of this amino acid contribute to salinity stress response in maize (Figure 1J). Collectively, these results demonstrate that NC326 and C68 exhibit distinct responses in tissue accumulation, physiological changes, and growth under salt stress, with NC326 showing greater resilience through better ion homeostasis and higher proline accumulation.

**Figure 1.**
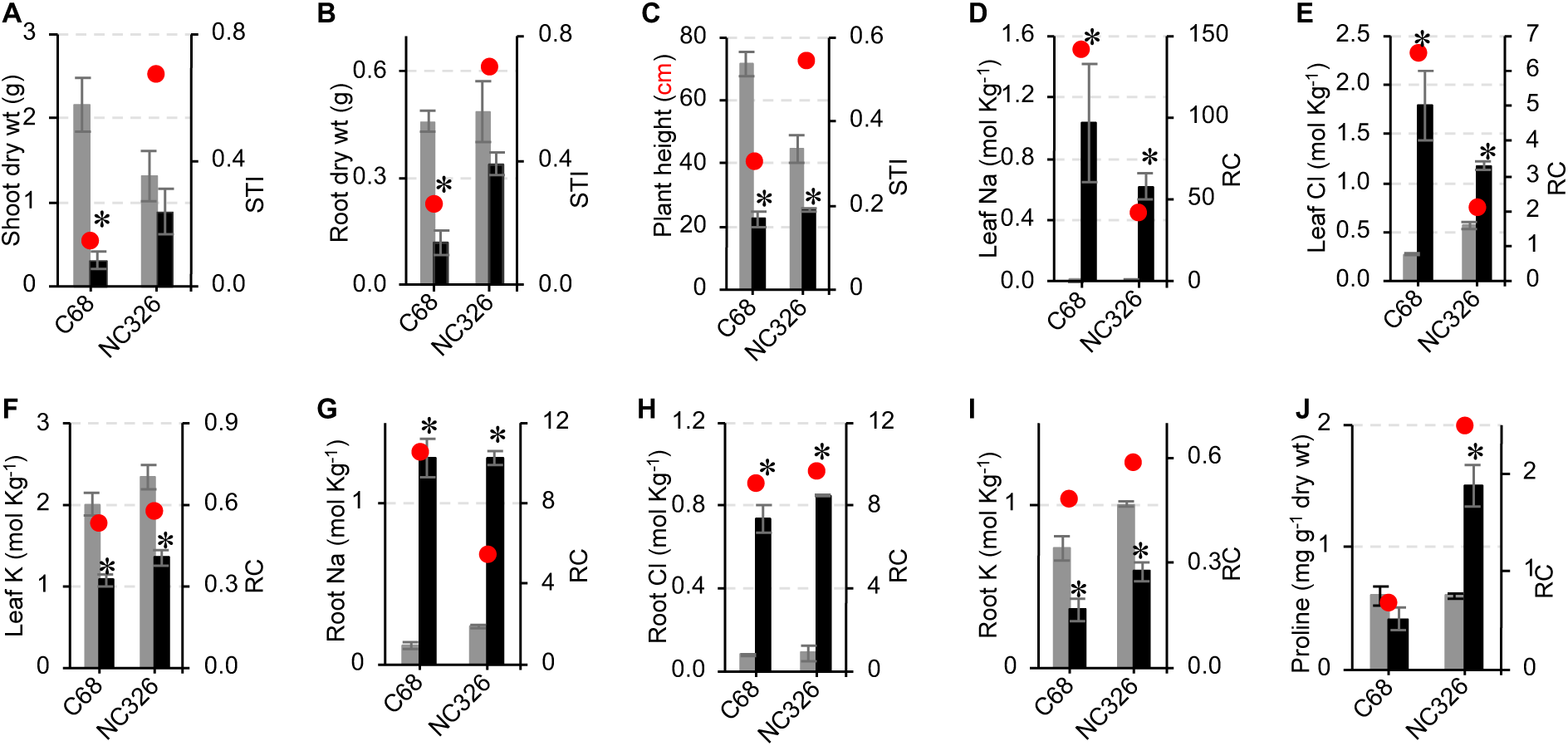
Salt stress impacts plant growth and physiology in maize. A-C, Effect of salinity on growth parameters including shoot dry weight (A), root dry weight (B), and plant height (C) with salt tolerance index (STI) shown by red circles. D-I, Effect of salinity on ion accumulation in the leaves (D-F) and roots (G-I). J, Effect of salinity on proline accumulation in leaves. Relative change (RC) is marked by red circles in D-J. Asterisks indicate significant differences between control (gray) and salinity (black) treatments (P ≤ 0.05, n = 3). Error bars represent the standard error.

### Global change in leaf and root metabolome in response to salt stress

To capture the metabolome variation underlying salt tolerance, we examined metabolite changes in the leaves and roots of NC326 and C68 inbred lines under control and salinity conditions. Untargeted metabolomic analysis detected 3,498 and 3,044 mass features in the leaf and root tissue, respectively. PCA based on leaf mass features revealed that PC1 and PC2 explained 31.6% and 24.1% variation in the metabolome, respectively (Figure 2A). Similarly, PCA based on root mass features showed that PC1 and PC2 explained 29.3% and 19.7 % variation in the root metabolome, respectively (Figure 2B). Remarkably, separate clustering of C68 and NC326 inbreds under control conditions in leaf and roots indicated that these inbred lines possess inherently distinct metabolic profiles. Furthermore, the leaf and root metabolome of the salt-sensitive inbred C68 had a clear distinction between salt stress and control conditions, indicating large-scale changes in response to salinity. However, in the salt-tolerant inbred NC326, we observed a less pronounced global distinction in leaf and root metabolomes between salt stress and control conditions, suggesting that targeted changes in specific metabolites, rather than broad metabolic shifts, are key to salinity tolerance.

**Figure 2.**
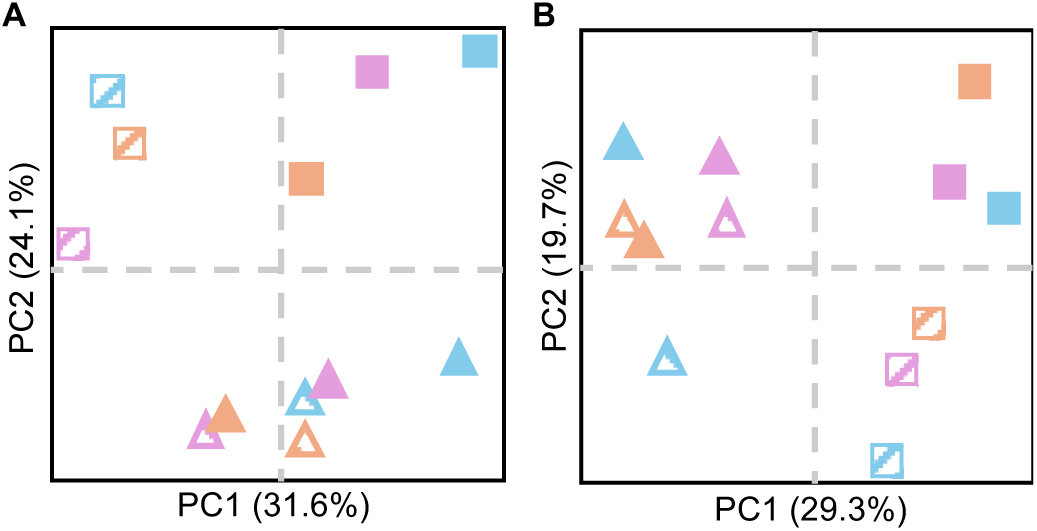
Salt stress drives temporal shifts in leaf and root metabolomes. Principal component analysis (PCA) illustrates temporal changes in the metabolome of leaves (A) and roots (B) during salt stress in salt-sensitive C68 (squares) and salt-tolerant NC326 (triangles) inbred lines. Each values represents the mean of three biological replications. Different colors represent distinct time points: 0 hr (sky blue), 24 hr (light purple), and 48 hr (light orange). Solid and pattern-filled shapes represent control and salinity treatment, respectively, and PC1 and PC2 denote the first and second principal components, respectively.

The lack of annotation of most identified mass features in metabolomic studies remains a major constraint due to the lack of reference spectra, preventing their designation as metabolites (Alseekh et al., 2021; da Silva et al., 2015). Of 3,498 mass features identified in leaves, we annotated 177 (5.06%) mass features as specialized metabolites belonging to 43 classes of organic compounds (Supplementary Dataset S1). Similarly, 143 (4.70%) of 3,044 mass features identified in the roots were annotated as specialized metabolites belonging to 51 classes of organic compounds (Supplementary Dataset S2). Phosphatidylcholines was the most abundant compound class, accounting for 17 metabolites each in leaf and root tissues indicating that these phospholipids play a vital role in salinity tolerance likely through improving in membrane stability and facilitating stress signaling (Guo et al., 2019).

### Constitutive accumulation of certain metabolites suggests adaptive priming

Since the global metabolome suggested inherent differences in metabolic profiles of the two inbred lines, we first compared the metabolomes of both inbred lines under control conditions to identify the baseline differences in metabolite accumulation. In leaves, 19 (10.7%) of the annotated specialized metabolites showed differential abundance between the two inbred lines (Supplementary Dataset S1), with 12 metabolites classified as flavonoids and the remaining seven representing unique compound classes. Of these, 11 flavonoids had significantly higher accumulation in the leaves of NC326 compared to C68 at all three timepoints (Figure 3, A-K). These include antioxidant flavonoids such as schaftoside, tricin, and kaempferol-related compounds that are expected to impart protection against excessive oxidative stress triggered by biotic or abiotic stresses. In roots, 11 (7.7%) of the annotated specialized metabolites differentially accumulated between the two inbred lines and, unlike leaves, all these metabolites belonged to unique classes of compounds (Supplementary Dataset S2). Roots of NC326 had constitutively high levels of a polyunsaturated fatty acid 28:6 at all three timepoints (Figure 3L). Constitutively high levels of certain specialized metabolites in leaves and roots of NC326 indicate that a fraction of the metabolome may be responsible for adaptive priming against salinity stress in maize. To ask how exposure to salt stress impacted these constitutive metabolites, we compared leaf and root tissues from the control and saline treatment. Because plants were gradually exposed to salt until reaching the desired water-salinity, achieved at 0h stage, we observed small differences in metabolite abundance between the C68 and NC326 inbred lines under both control and salinity conditions at 0h in both tissues (Figures 3). All the compounds maintained higher levels in NC326 compared to C68 under salt stress at all stages.

**Figure 3.**
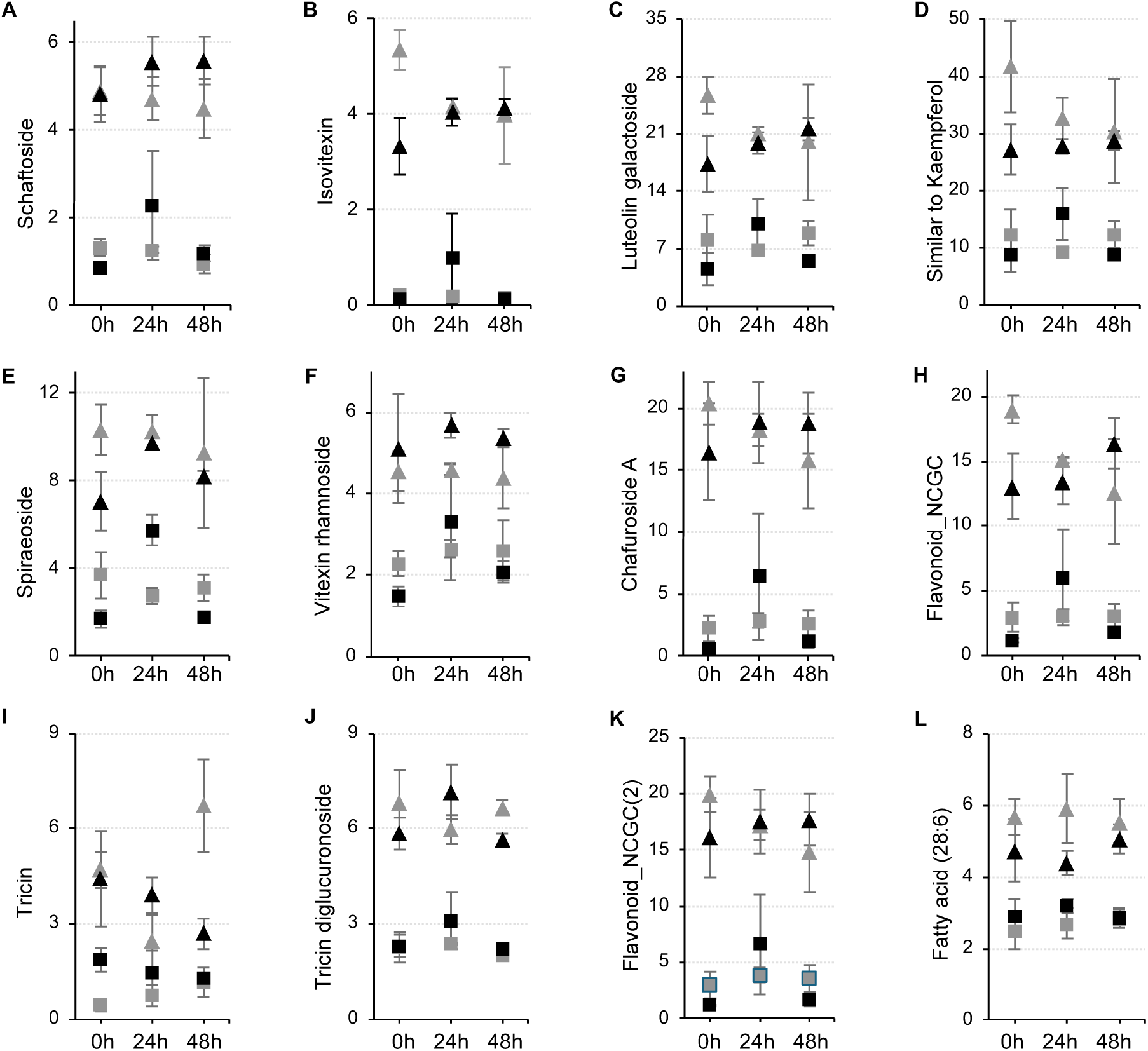
Adaptive priming of the metabolome underlies salt tolerance in maize. Metabolite levels in salt-tolerant NC326 (triangles) and salt-sensitive C68 (squares) are shown under control (gray) and salinity (black) conditions at different time points. (A-K), changes in various flavonoids in these inbred lines under control and salinity conditions. (L), levels of fatty acid (28:6) in roots.

### Identification of salt-responsive metabolites

To identify metabolites associated with salt stress, we compared the metabolome of both inbreds under control and salinity conditions for all three time-points. In leaves, 11 metabolites showed differential abundance in C68, while only three were differentially abundant in NC326 (Supplementary Dataset S1), suggesting that susceptible genotypes undergo broader metabolic shifts in response to salinity, whereas tolerant genotypes maintain a more stable metabolic profile. The majority (eight) of the differentially accumulated metabolites in C68 were upregulated. While most differentially abundant metabolites in leaves were specific to each inbred line, a few − including a galactolipid (DGDG;18:3/20:3) and a sterol (lanosterol) − were shared between both inbreds and exhibited similar changes in response to salt stress (Figure 4, B-C) (Supplementary Dataset S1). Consistent with the role of sterols in enhancing membrane stability (Du et al., 2022), lanosterol levels significantly increased in both inbred lines at all timepoints under salt stress. In contrast, galactolipid levels decreased in both inbreds at 0h, suggesting the reprogramming of plastid membranes in response to salt stress.

**Figure 4.**
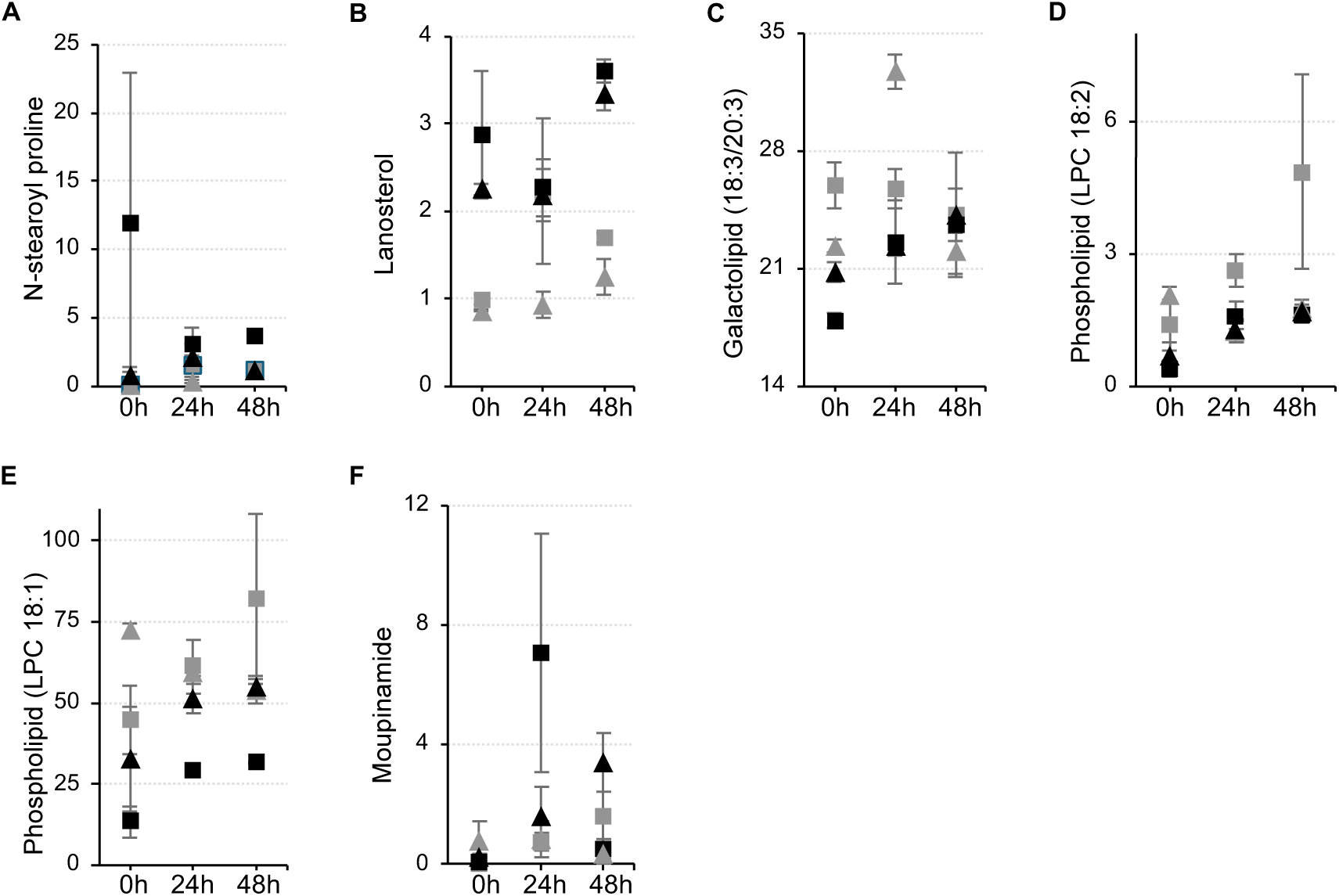
Metabolic changes associated with differential response to salinity. Levels of key metabolites in salt-tolerant NC326 (triangles) and salt-sensitive C68 (squares) are shown under control (gray) and salinity (black) conditions at different time points in leaves (A-C) and roots (D-F).

Similarly, in roots, seven metabolites were differentially accumulated in C68 while only three showed differential accumulation in NC326 inbred lines (Supplementary Dataset S2). About two-third (71%) of differentially abundant metabolites were phospholipids and downregulated in C68 roots, and two of these phospholipids downregulated in NC326 roots. Notably, two common lyso-phospholipids (LPC 18:1 and LPC 18:2) declined at all timepoints under salt stress in C68 roots, whereas in NC326, the levels of these metabolites decreased at 0h but later stabilized to levels comparable to control plants (Figure 4, D-E). This pattern of root metabolites maintaining normal levels after fluctuations under salinity in NC326 is consistent with observations of global metabolome and suggests that NC326 roots adapt more rapidly to salt stress, (Figure 2B). While the flux of phospholipids changed similarly in these inbreds, enhanced accumulation of metabolites with antioxidant properties was unique to C68 under salt stress. For example, moupinamide (also known as feruloyltyramine) specifically increased in C68 at 24h under salt stress, likely as a stress-mitigation response to oxidative damage (Figure 4F). These findings highlight distinct metabolic strategies between the inbreds, with NC326 showing early metabolic adjustments that may contribute to its salt tolerance.

### Identification of candidate genes underlying salt tolerance

Metabolites serve as functional readout of cellular biology and, therefore, provide a direct link between the genome and the external plant phenotypes. To understand the genetic architecture of salt tolerance, we identified the candidate genes involved in the biosynthesis of metabolites associated with response to salt stress. Our analyses revealed ten genes associated with eight metabolites, including eight genes linked to leaf metabolites and two to root metabolites (Table 2). Among these, UDP-glycosyltransferase, O-methyltransferase, and flavonol synthase were associated with the synthesis of antioxidant flavonoids such as schaftoside, tricin, and kaempferol-related compounds likely conferring stress protection through antioxidant activity or anti-inflammatory capacities. Genes encoding patatin-like phospholipase A, associated with phospholipid metabolism (LPC 18:3), likely mediate lipid remodeling to stabilize membranes under salinity. Additionally, N-hydroxycinnamoyl transferase, implicated in coumaroyl tyramine biosynthesis, might contribute to stress responses through phenolic metabolism. Besides, the two flavanol synthases and an O-methyltransferase, the role of the remaining eight genes in salt tolerance remains unexplored, suggesting that these are novel targets for improving in salt tolerance. Collectively, these candidate genes provide potential targets for improving maize salt tolerance through breeding or genetic engineering.

**Table 2:** Putative genes encoding enzymes upstream of metabolites.

| Metabolites | Enzymes | Putative genes |
| --- | --- | --- |
| Schaftoside | UDP-glycosyltransferase 708A6 | Zm00001d037382 |
| Tricin | O-methyltransferase | Zm00001d049541 |
| Similar to Kaempferol | Flavonol synthase | Zm00001d018187, Zm00001d018181 |
| Lanosterol | Lanosterol synthase | Zm00001d008674 |
| Amyrin | Terpene cyclase | Zm00001d052971, Zm00001d014695,<br>Zm00001d035389 |
| Isovitexin | C-glucosyl transferase | Zm00001d037382 |
| Coumaroyl tyramine | N-hydroxycinnamoyl transferase | Zm00001d048407 |
| LPC 18:3 | Patatin-like phospholipase A | Zm00001d031811 |

## Discussion

Salinity negatively impacts plant growth and vigor across all species, although salt-sensitive genotypes exhibit a more pronounced decline, suggesting the development of adaptive mechanisms to cope with salt stress. In this study, the salt-sensitive C68 inbred line showed a pronounced decrease in shoot dry weight, root dry weight, and plant height under salt stress compared to salt-tolerant NC326 inbred. While the concentration of Na and Cl increased significantly in the leaves of both inbred lines under salt stress, their relative accumulation of these salt ions and the resulting ion imbalance was significantly higher in C68. In contrast, both inbred lines had higher accumulation of these two ions in roots. These observations indicate that, compared to roots, ion imbalance in the leaves is more detrimental to plant growth and vigor likely due to disruption of photosynthesis, water balance, and other physiological functions. Additionally, NC326 maintained better Na⁺ homeostasis thus reducing the negative effects of excessive Na⁺ accumulation. Significantly higher proline in the leaves of NC326 indicates that the genetic differences for the synthesis of this osmoprotectant play a critical role in salt tolerance by stabilizing cellular functions, mitigating oxidative stress, and aiding in osmotic adjustment. Taken together, these results reinforce the findings from previous studies (El Moukhtari et al., 2020; Gupta & Huang, 2014; Sandhu et al., 2021) and underscore ion homeostasis and metabolic adjustments represent key physiological mechanisms conferring salt tolerance in maize.

Identification of specific metabolites differentiating salt-sensitive and salt-tolerant inbred lines is crucial for determining the metabolic pathways contributing to salt tolerance in maize. While the untargeted approach identified over 3,000 mass features each in root and leaves, only fewer than 5% of these features could be annotated to specific metabolites due to various reasons including the lack of reference spectra and insufficient published information (Alseekh et al., 2021). This significant gap in specialized metabolites annotation, consistent with our previous observations (Brar et al., 2025), poses a major challenge for uncovering mechanism and genes underlying salinity tolerance. However, this limitation presents exciting opportunities to characterize these unknown metabolites, which are significantly and differentially regulated between inbred lines and by salt stress. Future research including improved analytical techniques, metabolite (and unknown feature) association analyses for comprehensive linking of metabolites to salt tolerance, and metabolite (and unknown feature) quantitative trait locus (mQTL) mapping to identify genes and enzymes responsible for synthesizing key metabolites associated with salt tolerance will propel our knowledge of salt tolerance in maize and facilitate crop improvement strategies.

Metabolomics analysis revealed constitutively higher flavonoids in the leaves of NC326 compared to C68 under control conditions, suggesting that adaptive priming through enhanced accumulation of antioxidants including schaftoside, tricin, and kaempferol-related compounds could be a key mechanism underlying salt tolerance. Similar constitutive patterns were also observed in roots, although involving fewer metabolites primarily representing fatty acids, indicating that maintaining membrane fluidity and permeability may be the principal adaptive strategy employed by roots. For instance, schaftoside exhibits anti-inflammatory, antioxidant, and antiviral effects and is found in plants, including those used in traditional Chinese medicine (Lu et al., 2024). Schaftoside, the major bioactive compound in *Clinacanthus nutans* was shown to attenuate acute liver injury by inhibiting ferroptosis (cell death caused by iron-dependent lipid peroxidation) through the activation of the Nrf2/GPX4 (transcription factor/Glutathione peroxidase 4) pathway that is activated to protect cells from excessive reactive oxygen species. Tricin functions as a lignin monomer, providing support and rigidity to plant cell walls, especially in monocots, and is involved in plant growth, stress tolerance, and defense responses to pests (Li et al., 2016). Our results for tricin at 0h and 48h, although with no significant change at 24h, match the ones reported by Moheb et al. (2013) in that tricin concentrations tended to decrease in response to salt and drought stress compared to control plants at the same developmental stage. Thus, the expression of the flavonoids characterized here in response to salinity stress in corn plants have also found to play important roles in protecting human cells from damage caused by toxins and excessive reactive oxygen species. Although secondary metabolites from plants and animal cells are known to exert different roles in each respective organism, these flavonoids may have similar functions in both animal and plant cells as they protect both from cell-membrane and excessive ROS damage.

This constitutive metabolite upregulation could reflect inherent genetic differences arising from natural selection, or a correlated response to artificial selection for other stresses, given the limited targeted breeding for salinity stress in maize. Additionally, such adaptive priming might result from trans-generational epigenetic memory generated by past exposures of these genotypes to salt stress. Future systems genetics studies integrating genomic and epigenomic analyses will be valuable for disentangling the genetic and epigenetic contributions to salt tolerance. Furthermore, these studies may provide insights into potential resource allocation trade-offs and reveal the energetic costs associated with constitutive defense mechanisms and their impact on crop yields under non-stress conditions.

Compared to the constitutive metabolome, relatively fewer metabolites showed differential abundance in response to salinity suggesting that inducible changes in metabolome are more targeted. Higher number of metabolites responded to salt in C68 leaves and roots compared to NC326 suggesting that a large proportion of the inducible metabolome is involved in stress mitigation rather than prevention. For example, the metabolite moupinamide (feruloyltyramine) specifically increased in C68 at 24h under salt stress (Figure 4F), suggesting a potential role in mitigating oxidative damage. Feruloyltyramine has previously been shown to possess antioxidant properties and was reported to accumulate by up to ten-fold in tomato leaves following wounding (Pearce et al., 1998). Increased accumulation of antioxidant metabolites in C68 leaves and roots underscore the importance of certain metabolites in oxidative stress management. Besides genotype-specific responses, several metabolites were commonly responsive in both lines, reflecting shared mechanisms against salinity stress. For instance, increased lanosterol accumulation in leaves of both inbreds highlights critical role of this terpene in enhancing membrane integrity and decreasing ion permeability and transport (Chalbi et al., 2015; Kerkeb et al., 2001; Salama & Mansour, 2015). Lanosterol upregulation could also suggest an investment in the synthesis phytosterols (Ohyama et al., 2009) which could play a role in stress remediation. Conversely, the observed decrease in leaf galactolipids possibly reflects their conversion into sulfolipids, which enhance chloroplast membrane stability under salt stress (Guo et al., 2019). In roots, reduced lyso-phospholipid content under salt stress suggests changes in the lipid dynamics of the cells possibly strengthening membrane integrity with the goal of limiting ion leakage. Given that roots are the first line of defense against salt stress, phospholipid remodeling is a critical adaptive mechanism for maintaining membrane stability. Collectively, targeted adjustments in antioxidant production and membrane lipid composition are central to the inducible metabolic strategies that contribute to salt tolerance in maize.

The identification of 10 unique genes associated with the biosynthesis of eight specific metabolites provides valuable targets for improving salt tolerance in maize. The role of these candidate genes in conferring salt tolerance can be further validated through reverse genetic approaches. Seven genes involved in flavonoid biosynthesis are immediate candidates for crop improvement strategies. The identification of genes involved in altered metabolite regulation remain a significant limitation as only a few candidates could be identified in this study. To fully understand the genetic basis of complex traits, such as salt tolerance, future genomic approaches that include mQTL analyses and genome wide association studies are necessary

In conclusion, this study identified 56 metabolites underlying differential salt-stress responses in two maize inbred lines. Metabolomics revealed two key adaptive strategies including constitutive accumulation of flavonoids and lipids for antioxidant defense and membrane stabilization, and inducible metabolic adjustments targeting oxidative stress and membrane integrity. Increased salt induced lanosterol levels in both inbreds emphasized possible roles in maintaining membrane integrity, enhanced phytosterol synthesis and ion transport. Although 10 candidate genes regulating eight metabolites were identified as candidate targets for crop improvement, limited annotation restricted further genetic insights. Future studies employing mQTL mapping, metabolite-wide association analyses, and epigenomic approaches will enhance the genetic understanding of salt tolerance, informing targeted maize breeding strategies.

## Supporting information

List of identified mass features and differentially abundant metabolites in leaves of NC326 and C68 inbred lines under control and salinity conditions

List of identified mass features and differentially abundant metabolites in roots of NC326 and C68 inbred lines under control and salinity conditions.

## Supplementary Materials

Supplementary Dataset S1. List of identified mass features and differentially abundant metabolites in leaves of NC326 and C68 inbred lines under control and salinity conditions.

Supplementary Dataset S2. List of identified mass features and differentially abundant metabolites in roots of NC326 and C68 inbred lines under control and salinity conditions.

## Data availability

Data could be found in supplementary Dataset S1 (leaf metabolome) and S2 (root metabolome).

## Authorship contribution statement

**Manwinder S. Brar** – Methodology, Investigation, Data curation, Formal Analysis, Visualization, Writing – original draft, Writing – review & editing; **Amancio De Souza** - Methodology, Investigation, Data curation, Formal analysis, Writing – review & editing; **Avineet Ghai** – Investigation, Writing – review & editing; **Jorge F.S. Ferreira** – Formal analysis, Writing – review & editing; **Devinder Sandhu** – Conceptualization, Methodology, Investigation, Data curation, Formal analysis, Supervision, Resources, Funding acquisition, Project administration, Writing – original draft, Writing – review & editing; **Rajandeep S. Sekhon** – Conceptualization, Methodology, Investigation, Data curation, Formal analysis, Supervision, Resources, Funding acquisition, Project administration, Writing – original draft, Writing – review & editing.

## Acknowledgment

The authors thank Dr. Manju Pudussery, Dr. Xuan Liu, and Layton Chhour for technical help. This research was supported by USDA Agricultural Research Service project number 2036-13210-013-000D and National Science Foundation, Office of Integrative Activities award OIA# 1826715.

