## Supplementary material for "Untargeted metabolomics reveals key metabolites and genes underlying salinity tolerance mechanisms in maize": List of identified mass features and differentially abundant metabolites in leaves of NC326 and C68 inbred lines under control and salinity conditions

**Supplementary Dataset S1**: List of annotated mass features in leaves in NC326 and C68 inbred lines and differentially abundant (DA) metabolites under control and salinity conditions

| Sr No. | Final Accepted Description | Class | m/z | Retention time (min) | Annotation level | Compound | ClusterID | Neutral mass (Da) | Mass Error (ppm) | MS2Match | Adducts | Formula | LipidMAPS | DA in Control_C68 vs NC326 | DA in Salinity_C68 vs NC326 | DA in C68_Control vs Salinity | DA in NC326_Control vs Salinity |
|---|---|---|---|---|---|---|---|---|---|---|---|---|---|---|---|---|---|
| 1 | 10(E)_12(Z)-Conjugated Linoleic Acid | Fatty acid | 281.247639 | 17.68996667 | 1a | 17.69_281.2476m/z | 4 | | 0.581035398 | In-house | M+H | C18H32O2 | | - | - | X | - |
| 2 | N-stearoyl proline | Fatty Acyls | 364.320814 | 10.75103333 | 2b | 10.75_364.3208m/z | 794 | | -0.50438242 | Lipidmaps | M+H-H2O | C23H43NO3 | LMFA080201 19 | - | - | X | - |
| 3 | PC(18:3(9Z,12Z,15Z)/17:0) | PC | 770.568782 | 21.43585 | 2b | 21.44_770.5688m/z | 2008 | | -0.84374985 | Lipidmaps | M+H | C43H80NO8P | LMGP010116 79 | - | - | X | - |
| 4 | DGDG(18:3/20:3) | DGDG | 987.601142 | 20.80926667 | 2b | 20.81_964.6116n | 1820 | 964.611589 | -0.76013232 | Lipidblast | M+NH4, M+Na | C53H88O15 | | - | - | X | X |
| 5 | Triphenylphosphine oxide | Phosphine oxideÂ | 279.093434 | 22.82458333 | 2a | 22.82_279.0934m/z | 2303 | | 0.380066727 | Mona | M+H | C18H15OP | | - | - | X | X |
| 6 | Isorhamnetin | Flavonoid | 317.065382 | 7.098433333 | 2a | 7.10_317.0654m/z | 307 | | -0.62431049 | Mona | M+H | C16H12O7 | | - | X | - | - |
| 7 | Similar to Flavone base + 4O, C-Hex-dHex | Flavonoid glycoside | 595.16615 | 5.636633333 | 2a | 5.64_594.1589n | 141 | 594.158873 | 0.678085355 | Mona | M+H, M+Na | C27H30O15 | | - | X | - | - |
| 8 | Similar to Flavone base + 4O, C-Hex-dHex | Flavonoid glycoside | 595.166513 | 6.721833333 | 2a | 6.72_594.1592n | 138 | 594.159237 | 1.289854593 | Mona | M+H, M+Na | C27H30O15 | | - | X | - | - |
| 9 | PC(P-18:0/22:2(13Z,16Z)) | PC | 848.657784 | 23.71155 | 2b | 23.71_825.6686n | 223 | 825.668648 | 9.092667113 | Lipidmaps | M+H-H2O, M+H, M+Na | C48H92NO7P | LMGP010300 72 | - | X | - | - |
| 10 | PE-NMe(18:1(9Z)/16:0) | PE | 754.530883 | 21.32483333 | 2b | 21.32_754.5309m/z | 1981 | | -6.62051219 | Lipidmaps | M+Na | C40H78NO8P | LMGP020103 40 | - | X | - | - |
| 11 | PE(18:2(9Z,12Z)/17:0) | PE | 730.537465 | 21.71535 | 2b | 21.72_729.5302n | 2071 | 729.530188 | -0.91370933 | Lipidmaps | M+H, M+Na | C40H76NO8P | LMGP020106 60 | - | X | - | - |
| 12 | Similar to Campesterol | Sterol | 383.367566 | 22.22786667 | 2a | 22.23_383.3676m/z | 2168 | | 0.843389328 | Metlin | M+H-H2O | C28H48O | | - | X | - | - |
| 13 | Cholesterol | Sterol | 369.351715 | 21.7814 | 1a | 21.78_369.3517m/z | 2083 | | 0.353590475 | In-house | M+H-H2O | C27H46O | | - | X | - | - |
| 14 | Similar to Campesterol | Sterols | 383.367327 | 18.36993333 | 2a | 18.37_383.3673m/z | 1365 | | 0.246475925 | Metlin | M+H-H2O | C28H48O | | - | X | - | - |
| 15 | DGDG(16:0/18:3) | DGDG | 937.586034 | 20.89428333 | 2b | 20.89_937.5860m/z | 199 | | 0.154531053 | Lipidblast | M+Na | C49H86O15 | | - | X | X | - |
| 16 | Similar to Ricinoleic acid methyl ester | FAME | 313.274073 | 20.92703333 | 2a | 20.93_313.2741m/z | 1857 | | 1.125035841 | Metlin | M+H | C19H36O3 | | - | X | X | - |
| 17 | Similar to Ricinoleic acid methyl ester | FAME | 313.273937 | 22.1764 | 2a | 22.18_312.2667n | 2161 | 312.266661 | 0.691716827 | Metlin | M+H, M+Na | C19H36O3 | | - | X | X | - |
| 18 | Keioside | Flavonoid | 625.176684 | 6.766166667 | 1a | 6.77_625.1767m/z | 137 | | 0.652799818 | In-house | M+H | C28H32O16 | | - | X | X | - |
| 19 | ABOA | Carbohydrate conjugatesÂ | 178.050129 | 5.255833333 | 2a | 5.26_178.0501m/z | 126 | | 1.463464622 | Mona | M+H | C9H7NO3 | | X | - | - | - |
| 20 | Amyrin | Triterpenoid | 409.38251 | 22.7599 | 2a | 22.76_409.3825m/z | 2280 | | -0.86305471 | Metlin | M+H-H2O | C30H50O | | X | - | - | - |
| 21 | Tricin | Flavonoid | 331.081388 | 9.211033333 | 2a | 9.21_331.0814m/z | 750 | | 0.480907888 | Mona | M+H | C17H14O7 | | X | - | X | - |
| 22 | Lanosterol | Sterol | 409.382468 | 22.58566667 | 1a | 22.59_409.3825m/z | 2232 | | -1.03967344 | In-house | M+H-H2O | C30H50O | | X | - | X | X |
| 23 | Schaftoside | Flavonoid | 565.155557 | 5.707733333 | 2a | 5.71_565.1556m/z | 311 | | 0.664543677 | Mona | M+H | C26H28O14 | | X | X | - | - |
| 24 | Similar to Kaempferol | Flavonoid | 287.055404 | 7.038633333 | 2a | 7.04_287.0554m/z | 37 | | 1.362718831 | Mona | M+H | C15H10O6 | | X | X | - | - |
| 25 | Similar to Kaempferol | Flavonoid | 287.055158 | 9.50165 | 2a | 9.50_287.0552m/z | 757 | | 0.500222668 | Mona | M+H | C15H10O6 | | X | X | - | - |
| 26 | Chrysin 6-C-glucoside 8-C-arabinoside | Flavonoid glycoside | 549.160763 | 6.39815 | 2a | 6.40_549.1608m/z | 300 | | 0.90462202 | Mona | M+H | C26H28O13 | | X | X | - | - |
| 27 | chafuroside A | Flavonoid glycoside | 415.102276 | 7.484866667 | 2a | 7.48_414.0947n | 134 | 414.094719 | -0.87751942 | Mona | M+H-H2O, M+H | C21H18O9 | | X | X | - | - |
| 28 | Tricin diglucuronoside | Flavonoid glycoside | 683.145196 | 6.953733333 | 2a | 6.95_683.1452m/z | 668 | | -0.30739554 | Mona | M+H | C29H30O19 | | X | X | - | - |
| 29 | Isovitexin(4) | Flavonoid glycoside | 433.112953 | 6.34265 | 2a | 6.34_432.1054n | 304 | 432.105419 | -0.5274342 | Mona | M+H-H2O, M+H | C21H20O10 | | X | X | - | - |
| 30 | Spiraeoside | Flavonoid glycoside | 487.08481 | 6.625 | 2a | 6.63_464.0959n | 8 | 464.09594 | 0.999233784 | Mona | M+H, M+Na | C21H20O12 | | X | X | - | - |
| 31 | Luteolin 7-galactoside | Flavonoid glycoside | 471.090102 | 7.036316667 | 1a | 7.04_448.1008n | 677 | 448.100829 | -0.38131796 | In-house | M+H, M+Na | C21H20O11 | | X | X | - | - |
| 32 | Similar to Vitexin-2-rhamnoside | Flavonoid glycoside | 579.171388 | 6.129366667 | 2a | 6.13_579.1714m/z | 303 | | 0.961819072 | Mona | M+H | C27H30O14 | | X | X | - | - |
| 33 | Similar to Vitexin-2-rhamnoside | Flavonoid glycoside | 579.171177 | 7.266983333 | 2a | 7.27_579.1712m/z | 80 | | 0.596028116 | Mona | M+H | C27H30O14 | | X | X | - | - |
| 34 | Glucocerebrosides | Glucocerebroside | 696.537133 | 20.0155 | 2a | 20.02_713.5435n | 49 | 713.543518 | -0.93155613 | Metlin | M+H-H2O, M+H, M+Na | C40H75NO9 | | X | X | - | - |
| 35 | Citroside A | Terpene glycoside | 409.182468 | 5.37445 | 1a | 5.37_409.1825m/z | 66 | | -1.94891197 | In-house | M+Na | C19H30O8 | | X | X | - | - |
| 36 | Similar to CHLOROGENIC ACID | Organooxygen compounds | 377.084379 | 4.940866667 | 2a | 4.94_377.0844m/z | 75 | | 0.213892711 | Mona | M+Na | C16H18O9 | | X | X | X | - |
| 37 | Similar to CHLOROGENIC ACID | Organooxygen compounds | 377.084479 | 5.557166667 | 2a | 5.56_377.0845m/z | 140 | | 0.49617218 | Mona | M+Na | C16H18O9 | | X | X | X | - |
| 38 | CETRIMONIUM | Amine | 284.33128 | 11.30608333 | 2a | 11.31_284.3313m/z | 812 | | 0.362559224 | Metlin | M+H | C19H41N | | - | - | - | - |
| 39 | SPHINGANINE | Amine | 302.305453 | 10.2562 | 2a | 10.26_302.3055m/z | 782 | | 0.320732025 | Mona | M+H | C18H39NO2 | | - | - | - | - |
| 40 | Tetradecyldiethanolamine | Amine | 302.305305 | 9.520816667 | 2a | 9.52_302.3053m/z | 760 | | -0.17084152 | Mona | M+H | C18H39NO2 | | - | - | - | - |
| 41 | Dihydroactinidiolide | Benzofurans | 181.122544 | 9.588216667 | 2a | 9.59_181.1225m/z | 762 | | 1.317547564 | Metlin | M+H | C11H16O2 | | - | - | - | - |
| 42 | Loliolide | Benzofurans | 197.117448 | 6.744816667 | 2a | 6.74_196.1100n | 649 | 196.109957 | 0.062696496 | Mona | M+H-H2O, M+H | C11H16O3 | | - | - | - | - |
| 43 | HMBOA-Glc | Carbohydrate conjugatesÂ | 380.095563 | 5.244333333 | 1a | 5.24_380.0956m/z | 68 | | 0.957669384 | In-house | M+Na | C15H19NO9 | | - | - | - | - |
| 44 | Pheophorbide A | Chlorophyll breakdown | 593.276466 | 18.58903333 | 2a | 18.59_592.2692n | 87 | 592.269189 | 1.045225789 | Mona | M+H, M+Na | C35H36N4O5 | | - | - | - | - |
| 45 | Coumaric acid | Cinnamic acids and derivatives | 147.044363 | 7.313033333 | 2a | 7.31_164.0477n | 130 | 164.047651 | 1.870962584 | Mona | M+H-H2O, M+H | C9H8O3 | | - | - | - | - |
| 46 | Coumarin | Coumarins and derivativesÂ | 147.044365 | 5.504233333 | 2a | 5.50_147.0444m/z | 312 | | 2.119268784 | Metlin | M+H | C9H6O2 | | - | - | - | - |
| 47 | DG(16:0/16:1(9Z)/0:0)[iso2] | DG | 589.481476 | 19.90475 | 2b | 19.90_589.4815m/z | 1597 | | 2.170916498 | Lipidmaps | M+Na | C35H66O5 | LMGL020100 10 | - | - | - | - |
| 48 | Similar to DG(16:0/18:3(9Z,12Z,15Z)/0:0)[iso2] | DG | 591.498973 | 20.92463333 | 2b | 20.92_590.4907n | 200 | 590.490704 | -0.5434797 | Lipidmaps | M+H-H2O, M+H | C37H66O5 | LMGL020100 32 | - | - | - | - |

| | | | | | | | | | | | | | | | | | |
|---|---|---|---|---|---|---|---|---|---|---|---|---|---|---|---|---|---|
| 49 | Similar to DG(16:0/18:3(9Z,12Z,15Z)/0:0)[iso2] | DG | 573.487492 | 22.1764 | 2b | 22.18_590.4908n | 2162 | 590.49078 | -0.4150041 | Lipidmaps | M+H-H2O, M+H | C37H66O5 | LMGL02010032 | - | - | - | - |
| 50 | DG(16:1(9Z)/18:3(9Z,12Z,15Z)/0:0)[iso2] | DG | 571.472154 | 20.8672 | 2b | 20.87_571.4722m/z | 1838 | | 0.113721043 | Lipidmaps | M+H-H2O | C37H64O5 | LMGL02010035 | - | - | - | - |
| 51 | DG(18:0/18:2(9Z,12Z)/0:0)[iso2] | DG | 603.534067 | 22.4469 | 2b | 22.45_603.5341m/z | 2208 | | -0.99960654 | Lipidmaps | M+H-H2O | C39H72O5 | LMGL02010050 | - | - | - | - |
| 52 | DG(18:0/18:3(9Z,12Z,15Z)/0:0)[iso2] | DG | 619.52957 | 21.90156667 | 2b | 21.90_618.5212n | 201 | 618.521207 | -1.80783026 | Lipidmaps | M+H-H2O, M+H | C39H70O5 | LMGL02010057 | - | - | - | - |
| 53 | DG(16:0/20:4(5Z,8Z,11Z,14Z)/0:0)[iso2] | DG | 639.495297 | 23.7047 | 2b | 23.70_616.5065n | 219 | 616.50655 | -0.20355374 | Lipidmaps | M+H, M+Na | C39H68O5 | LMGL02010070 | - | - | - | - |
| 54 | DG(12:0/22:5(7Z,10Z,13Z,16Z,19Z)/0:0)[iso2] | DG | 609.450946 | 16.39208333 | 2b | 16.39_609.4509m/z | 93 | | 3.411297294 | Lipidmaps | M+Na | C37H62O5 | LMGL02010352 | - | - | - | - |
| 55 | DG(14:1(9Z)/20:3(8Z,11Z,14Z)/0:0)[iso2] | DG | 571.471857 | 21.29146667 | 2b | 21.29_588.4751n | 1973 | 588.475145 | -0.39104794 | Lipidmaps | M+H-H2O, M+H | C37H64O5 | LMGL02010415 | - | - | - | - |
| 56 | DG(18:3(6Z,9Z,12Z)/18:3(9Z,12Z,15Z)/0:0)[iso2] | DG | 635.464001 | 22.85243333 | 2b | 22.85_612.4761n | 2311 | 612.476086 | 1.160477689 | Lipidmaps | M+NH4, M+Na | C39H64O5 | LMGL02010482 | - | - | - | - |
| 57 | DIPALMITOYLGLYCEROL | DG | 551.497889 | 21.29146667 | 2a | 21.29_551.4979m/z | 1971 | | -9.71423654 | In-house | M+H-H2O | C35H68O5 | | - | - | - | - |
| 58 | DGDG(16:0/16:0) | DGDG | 915.600373 | 21.92231667 | 2b | 21.92_892.6111n | 2119 | 892.611136 | -1.32877315 | Lipidblast | M+NH4, M+Na | C47H88O15 | | - | - | - | - |
| 59 | DGDG(16:0/18:1) | DGDG | 941.616292 | 22.16601667 | 2b | 22.17_918.6264n | 2160 | 918.626416 | -1.69424929 | Lipidblast | M+NH4, M+Na | C49H90O15 | | - | - | - | - |
| 60 | DGDG(16:0/18:2) | DGDG | 939.600894 | 21.48895 | 2b | 21.49_939.6009m/z | 2024 | | -0.70805173 | Lipidblast | M+Na | C49H88O15 | | - | - | - | - |
| 61 | DGDG(16:1/18:3) | DGDG | 935.569572 | 20.18773333 | 2b | 20.19_912.5804n | 1653 | 912.580356 | -0.73002141 | Lipidblast | M+NH4, M+Na | C49H84O15 | | - | - | - | - |
| 62 | DGDG(17:0/18:3) | DGDG | 951.600604 | 21.4182 | 2b | 21.42_928.6116n | 2004 | 928.611582 | -0.79660954 | Lipidblast | M+NH4, M+Na | C50H88O15 | | - | - | - | - |
| 63 | DGDG(2:0/8:0) | DGDG | 607.254923 | 25.16528333 | 2b | 25.17_607.2549m/z | 2623 | | -3.96878501 | Lipidblast | M+Na | C25H44O15 | | - | - | - | - |
| 64 | Methyl linolenate | FAME | 293.247512 | 18.78286667 | 2a | 18.78_293.2475m/z | 1413 | | 0.017414758 | Metlin | M+H | C19H32O2 | | - | - | - | - |
| 65 | formyl 2E,4E,6Z-decatrienoate | FAME | 181.122443 | 12.14801667 | 2b | 12.15_181.1224m/z | 831 | | 0.758265027 | Lipidmaps | M+H | C11H16O2 | LMFA07010481 | - | - | - | - |
| 66 | Similar to Linolenic Acid | Fatty acid | 279.232006 | 14.03291667 | 2a | 14.03_278.2246n | 266 | 278.224552 | -0.10008084 | Mona | M+H-H2O, M+H | C18H30O2 | | - | - | - | - |
| 67 | 5,8,11-dodecatriynoic acid | Fatty acid | 189.091825 | 24.9801 | 2b | 24.98_189.0918m/z | 2593 | | 4.351236813 | Lipidmaps | M+H | C12H12O2 | LMFA01030465 | - | - | - | - |
| 68 | 10-oxo-nonadecanoic acid | Fatty acid | 335.25823 | 21.90156667 | 2b | 21.90_335.2582m/z | 2114 | | 8.212293363 | Lipidmaps | M+Na | C19H36O3 | LMFA01060128 | - | - | - | - |
| 69 | 2-methyl-tridecanedioic acid | Fatty acid | 259.190561 | 20.10161667 | 2b | 20.10_258.1832n | 170 | 258.183247 | 0.534520681 | Lipidmaps | M+H-H2O, M+H | C14H26O4 | LMFA01170015 | - | - | - | - |
| 70 | (9S,13S)-10,11-dihydro-12-oxo-15-phytoenoic acid | Fatty acid | 277.216062 | 12.78505 | 2b | 12.79_277.2161m/z | 857 | | -0.49243267 | Lipidmaps | M+H-H2O | C18H30O3 | LMFA02010007 | - | - | - | - |
| 71 | Similar to Linolenic Acid | Fatty acid | 279.232246 | 16.8511 | 1a | 16.85_278.2248n | 1 | 278.224781 | 0.652098007 | In-house | M+H-H2O, M+H | C18H30O2 | | - | - | - | - |
| 72 | 12,14-octadecadiynoic acid | Fatty Acyls | 277.216452 | 13.23668333 | 2b | 13.24_276.2090n | 12 | 276.208972 | 0.153008221 | Lipidmaps | M+H-H2O, M+H | C18H28O2 | LMFA01030547 | - | - | - | - |
| 73 | 12,16-octadecadiynoic acid | Fatty Acyls | 277.216304 | 13.40638333 | 2b | 13.41_276.2090n | 263 | 276.208981 | 0.183969247 | Lipidmaps | M+H-H2O, M+H | C18H28O2 | LMFA01030549 | - | - | - | - |
| 74 | Juniperic acid | Fatty Acyls | 290.269014 | 8.630516667 | 2b | 8.63_290.2690m/z | 737 | | 0.161210446 | Lipidmaps | M+NH4 | C16H32O3 | LMFA01050051 | - | - | - | - |
| 75 | 15,16-EpODE | Fatty Acyls | 333.204027 | 11.70696667 | 2b | 11.71_310.2145n | 278 | 310.214515 | 0.338695744 | Lipidmaps | M+H-H2O, M+Na | C18H30O4 | LMFA02000030 | - | - | - | - |
| 76 | Similar to 17-hydroxy-linolenic acid | Fatty Acyls | 277.216126 | 12.60035 | 2b | 12.60_277.2161m/z | 849 | | -0.27557944 | Lipidmaps | M+H-H2O | C18H30O3 | LMFA02000239 | - | - | - | - |
| 77 | Similar to 17-hydroxy-linolenic acid | Fatty Acyls | 277.216157 | 14.36536667 | 2b | 14.37_277.2162m/z | 941 | | -0.16846885 | Lipidmaps | M+H-H2O | C18H30O3 | LMFA02000239 | - | - | - | - |
| 78 | 3-Methyl-3-butenyl hexadecanoate | Fatty Acyls | 342.336628 | 11.63996667 | 2b | 11.64_342.3366m/z | 821 | | -0.0860595 | Lipidmaps | M+NH4 | C21H40O2 | LMFA07010635 | - | - | - | - |
| 79 | Anandamide (18:3, n-3) | Fatty amide | 344.258446 | 8.00775 | 2b | 8.01_344.2584m/z | 723 | | 7.612388943 | Lipidmaps | M+Na | C20H35NO2 | LMFA08040007 | - | - | - | - |
| 80 | Quercetin | Flavonoid | 303.049817 | 8.571133333 | 1a | 8.57_303.0498m/z | 735 | | -0.52911023 | In-house | M+H | C15H10O7 | | - | - | - | - |
| 81 | Taxifolin | Flavonoid | 287.055508 | 6.721833333 | 2a | 6.72_304.0588n | 635 | 304.058797 | 2.61960578 | In-house | M+H-H2O, M+Na | C15H12O7 | | - | - | - | - |
| 82 | Similar to NCGC00384960-01!5,7-dihydroxy-6-[(2S,3R,4R,6R)-4-hydroxy-6-methyl-5-oxo-3-[(2S,3R,4R,5R,6S)-3,4,5-trihydroxy-6-methyloxan-2-yl]oxyoxan-2-yl]-2-(4-hydroxyphenyl)chromen-4-one [IIN-based: Match] | Flavonoid glycoside | 561.160637 | 7.493816667 | 2a | 7.49_561.1606m/z | 134 | | 0.660102076 | Mona | M+H | C27H28O13 | | - | - | - | - |
| 83 | Similar to NCGC00384960-01!5,7-dihydroxy-6-[(2S,3R,4R,6R)-4-hydroxy-6-methyl-5-oxo-3-[(2S,3R,4R,5R,6S)-3,4,5-trihydroxy-6-methyloxan-2-yl]oxyoxan-2-yl]-2-(4-hydroxyphenyl)chromen-4-one [IIN-based: Match] | Flavonoid glycoside | 561.160692 | 7.692666667 | 2a | 7.69_560.1534n | 21 | 560.153416 | 0.758354332 | Mona | M+H, M+Na | C27H28O13 | | - | - | - | - |
| 84 | Ilixathin | Flavonoid glycoside | 611.161391 | 6.328183333 | 1a | 6.33_610.1541n | 20 | 610.154115 | 1.82730231 | In-house | M+H, M+Na | C27H30O16 | | - | - | - | - |
| 85 | 2-Linoleoyl Glycerol | Glycerin fatty acid ester | 337.273456 | 16.2894 | 1a | 16.29_354.2767n | 94 | 354.276744 | -0.72231966 | In-house | M+H-H2O, M+H, M+Na | C21H38O4 | | - | - | - | - |
| 86 | Similar to NCGC00380867-01_C27H46O9_9,12,15-Octadecatrienoic acid, 3-(hexopyranosyloxy)-2-hydroxypropyl ester, (9Z,12Z,15Z)- | Glycoglycerolipid | 532.34851 | 13.38926667 | 2a | 13.39_532.3485m/z | 894 | | 0.975746093 | Mona | M+NH4 | C27H46O9 | | - | - | - | - |

| | | | | | | | | | | | | | | | | | |
|---|---|---|---|---|---|---|---|---|---|---|---|---|---|---|---|---|---|
| 87 | Similar to NCGC00380867-01_C27H46O9_9,12,15-Octadecatrienoic acid, 3-(hexopyranosyloxy)-2-hydroxypropyl ester, (9Z,12Z,15Z)- | Glycoglycerolipid | 537.304047 | 13.40638333 | 2a | 13.41_514.3145n | 32 | 514.314517 | 0.648960037 | Mona | M+H, M+Na | C27H46O9 | | - | - | - | - |
| 88 | Similar to Glc-GP(18:0/20:4(5Z,8Z,11Z,14Z)) | GP | 887.567171 | 24.84786667 | 2b | 24.85_886.5599n | 2579 | 886.559894 | 3.118757301 | Lipidmaps | M+H, M+Na | C47H83O13P | LMGP140100 01 | - | - | - | - |
| 89 | Similar to Glc-GP(18:0/20:4(5Z,8Z,11Z,14Z)) | GP | 869.55732 | 25.1262 | 2b | 25.13_869.5573m/z | 2619 | | 3.923498198 | Lipidmaps | M+H-H2O | C47H83O13P | LMGP140100 01 | - | - | - | - |
| 90 | Similar to Glc-GP(18:0/20:4(5Z,8Z,11Z,14Z)) | GP | 869.557175 | 25.2754 | 2b | 25.28_869.5572m/z | 2641 | | 3.759990737 | Lipidmaps | M+H-H2O | C47H83O13P | LMGP140100 01 | - | - | - | - |
| 91 | Similar to 3-methyl-1-heptene | Hydrocarbon | 130.159407 | 27.80093333 | 2b | 27.80_130.1594m/z | 2729 | | 3.394261536 | Lipidmaps | M+NH4 | C8H16 | LMFA110005 92 | - | - | - | - |
| 92 | PC(18:1(6Z)/0:0) | Lyso PC | 522.355631 | 13.70605 | 2b | 13.71_521.3484n | 911 | 521.348354 | 0.411497587 | Lipidmaps | M+H, M+Na | C26H52NO7P | LMGP010500 29 | - | - | - | - |
| 93 | PC(18:3(9Z,12Z,15Z)/0:0) | Lyso PC | 518.324418 | 12.19406667 | 2b | 12.19_517.3171n | 63 | 517.317141 | 0.583042802 | Lipidmaps | M+H, M+Na | C26H48NO7P | LMGP010500 38 | - | - | - | - |
| 94 | PC(16:0/0:0) | Lyso PC | 496.339966 | 13.42501667 | 2a | 13.43_495.3327n | 904 | 495.33269 | 0.403669488 | Mona | M+H, M+Na | C24H50NO7P | | - | - | - | - |
| 95 | PC(18:1/0:0) | Lyso PC | 522.355674 | 13.96576667 | 2b | 13.97_522.3557m/z | 925 | | 0.495270439 | Lipidblast | M+H | C26H52NO7P | | - | - | - | - |
| 96 | PC(18:2/0:0) | Lyso PC | 520.340333 | 13.1436 | 2b | 13.14_519.3331n | 257 | 519.333057 | 1.092142655 | Lipidblast | M+H-H2O, M+H, M+Na | C26H50NO7P | | - | - | - | - |
| 97 | PC(18:3/0:0) | Lyso PC | 518.324317 | 12.44446667 | 2b | 12.44_517.3170n | 843 | 517.31704 | 0.38770132 | Lipidblast | M+H, M+Na | C26H48NO7P | | - | - | - | - |
| 98 | PE(16:0/0:0) | Lyso PE | 454.292905 | 13.35436667 | 2a | 13.35_453.2856n | 891 | 453.285629 | 0.19685327 | Metlin | M+H-H2O, M+H, M+Na | C21H44NO7P | | - | - | - | - |
| 99 | PE(18:2(9Z,12Z)/0:0) | Lyso PE | 478.293297 | 12.81905 | 2b | 12.82_477.2860n | 269 | 477.28602 | 1.007319427 | Lipidmaps | M+H, M+Na | C23H44NO7P | LMGP020500 11 | - | - | - | - |
| 100 | PG(18:4(6Z,9Z,12Z,15Z)/0:0) | Lyso PG | 522.282803 | 15.7806 | 2b | 15.78_522.2828m/z | 1034 | | 0.313096858 | Lipidmaps | M+NH4 | C24H41O9P | LMGP040500 21 | - | - | - | - |
| 101 | PG(18:3(9Z,12Z,15Z)/0:0) | Lyso PG | 529.254087 | 18.75405 | 2b | 18.75_506.2651n | 158 | 506.265131 | 1.30706615 | Lipidmaps | M+H-H2O, M+H, M+Na | C24H43O9P | LMGP040500 32 | - | - | - | - |
| 102 | MGDG(20:5(5Z,8Z,11Z,14Z,17Z)/16:3(7Z,10Z,13Z)) | MGDG | 771.501772 | 21.01993333 | 2b | 21.02_771.5018m/z | 1881 | | -3.11891051 | Lipidmaps | M+H | C45H70O10 | LMGL050100 17 | - | - | - | - |
| 103 | MGDG(16:0/18:2(9Z,12Z)) | MGDG | 777.548738 | 22.69611667 | 2b | 22.70_777.5487m/z | 2273 | | 0.024621504 | Lipidmaps | M+Na | C43H78O10 | LMGL050100 26 | - | - | - | - |
| 104 | MGDG(18:1/16:0) | MGDG | 779.564149 | 23.25006667 | 2b | 23.25_779.5641m/z | 2385 | | -0.29086498 | Lipidblast | M+Na | C43H80O10 | | - | - | - | - |
| 105 | MGDG(18:1/18:2) | MGDG | 803.563345 | 22.9917 | 2b | 22.99_803.5633m/z | 2338 | | -1.31249994 | Lipidblast | M+Na | C45H80O10 | | - | - | - | - |
| 106 | MGDG(18:3/16:1) | MGDG | 773.516188 | 21.66878333 | 2b | 21.67_773.5162m/z | 2054 | | -1.64012897 | Lipidblast | M+Na | C43H74O10 | | - | - | - | - |
| 107 | MGDG(18:3/20:3) | MGDG | 825.550447 | 22.03221667 | 2b | 22.03_825.5504m/z | 238 | | 2.152041771 | Lipidblast | M+Na | C47H78O10 | | - | - | - | - |
| 108 | Similar to Lovastatin acid | Naphthalenes | 423.274023 | 13.3762 | 2a | 13.38_422.2666n | 260 | 422.266628 | -0.49998628 | Mona | M+H-H2O, M+H | C24H38O6 | | - | - | - | - |
| 109 | Similar to Lovastatin acid | Naphthalenes | 423.274273 | 21.01356667 | 2a | 21.01_422.2666n | 52 | 422.266568 | -0.64259283 | Mona | M+H-H2O, M+H | C24H38O6 | | - | - | - | - |
| 110 | N-Phenyl-2-naphthylamine | Naphthalenes | 220.11227 | 14.71638333 | 2a | 14.72_220.1123m/z | 972 | | 0.886516136 | Mona | M+H | C16H13N | | - | - | - | - |
| 111 | Similar to 9,11-methane-epoxy PGF1 | Oxylipin | 353.268686 | 12.01105 | 2a | 12.01_352.2615n | 61 | 352.261461 | 0.286512976 | Metlin | M+H-H2O, M+H | C21H36O4 | | - | - | - | - |
| 112 | Similar to 9,11-methane-epoxy PGF1 | Oxylipin | 353.268753 | 12.28405 | 2a | 12.28_352.2611n | 14 | 352.261101 | -0.73301674 | Metlin | M+H-H2O, M+H | C21H36O4 | | - | - | - | - |
| 113 | Similar to 9,11-methane-epoxy PGF1 | Oxylipin | 353.269094 | 13.16406667 | 2a | 13.16_352.2615n | 254 | 352.261519 | 0.451263158 | Metlin | M+H-H2O, M+H | C21H36O4 | | - | - | - | - |
| 114 | Similar to 9,11-methane-epoxy PGF1 | Oxylipin | 353.268945 | 13.40411667 | 2a | 13.40_352.2617n | 119 | 352.261735 | 1.064967916 | Metlin | M+H-H2O, M+H | C21H36O4 | | - | - | - | - |
| 115 | Similar to 9,11-methane-epoxy PGF1 | Oxylipin | 353.268489 | 15.45166667 | 2a | 15.45_352.2612n | 92 | 352.261157 | -0.5748893 | Metlin | M+H-H2O, M+H | C21H36O4 | | - | - | - | - |
| 116 | Similar to 9,11-methane-epoxy PGF1 | Oxylipin | 353.268497 | 21.00716667 | 2a | 21.01_353.2685m/z | 52 | | -0.39422363 | Metlin | M+H | C21H36O4 | | - | - | - | - |
| 117 | Similar to 13-epi-12-oxo Phytodienoic Acid | Oxylipin | 293.211142 | 12.91321667 | 1a | 12.91_292.2039n | 861 | 292.203885 | 0.290841111 | In-house | M+H-H2O, M+H, M+Na | C18H28O3 | | - | - | - | - |
| 118 | Similar to 13-epi-12-oxo Phytodienoic Acid | Oxylipin | 275.200487 | 13.87036667 | 2a | 13.87_292.2038n | 920 | 292.203775 | -0.08602114 | In-house | M+H-H2O, M+H, M+Na | C18H28O3 | | - | - | - | - |
| 119 | PC(18:3(6Z,9Z,12Z)/18:3(6Z,9Z,12Z)) | PC | 778.538126 | 19.91116667 | 2a | 19.91_778.5381m/z | 1603 | | -0.00706191 | Metlin | M+H | C44H76NO8P | | - | - | - | - |
| 120 | GPCho(16:0/18:3) | PC | 778.53576 | 20.97785 | 2b | 20.98_778.5358m/z | 1868 | | 0.044827529 | Lipidblast | M+Na | C42H78NO8P | | - | - | - | - |
| 121 | GPCho(18:3/18:2) | PC | 802.535621 | 20.56276667 | 2b | 20.56_802.5356m/z | 1767 | | -0.13496583 | Lipidblast | M+Na | C44H78NO8P | | - | - | - | - |
| 122 | GPCho(18:3/18:3) | PC | 800.520337 | 19.93195 | 2b | 19.93_800.5203m/z | 1614 | | 0.335666096 | Lipidblast | M+Na | C44H76NO8P | | - | - | - | - |
| 123 | PC(16:0/16:0) | PC | 734.568767 | 21.92723333 | 2b | 21.93_734.5688m/z | 2120 | | -0.90630303 | Lipidmaps | M+H | C40H80NO8P | LMGP010105 64 | - | - | - | - |
| 124 | PC(16:0/18:2(10E,12Z)) | PC | 758.569692 | 21.55685 | 2b | 21.56_758.5697m/z | 2029 | | 0.343177839 | Lipidmaps | M+H | C42H80NO8P | LMGP010105 85 | - | - | - | - |
| 125 | PC(18:2(2Z,4Z)/18:2(2Z,4Z)) | PC | 782.569405 | 21.15606667 | 2b | 21.16_781.5621n | 1919 | 781.562128 | -0.0344617 | Lipidmaps | M+H, M+Na | C44H80NO8P | LMGP010109 24 | - | - | - | - |
| 126 | PC(18:2(9Z,12Z)/20:1(13Z)) | PC | 812.615301 | 22.68068333 | 2b | 22.68_812.6153m/z | 2265 | | -1.33169883 | Lipidmaps | M+H | C46H86NO8P | LMGP010109 42 | - | - | - | - |
| 127 | PC(18:2(9Z,12Z)/14:0) | PC | 730.537516 | 20.56515 | 2b | 20.57_730.5375m/z | 1771 | | -0.84315153 | Lipidmaps | M+H | C40H76NO8P | LMGP010116 16 | - | - | - | - |
| 128 | PC(18:3(6Z,9Z,12Z)/15:0) | PC | 742.536757 | 20.44673333 | 2b | 20.45_742.5368m/z | 1747 | | -1.85413603 | Lipidmaps | M+H | C41H76NO8P | LMGP010116 44 | - | - | - | - |
| 129 | PC(18:3(6Z,9Z,12Z)/16:1(9Z)) | PC | 754.537367 | 20.32051667 | 2b | 20.32_754.5374m/z | 1660 | | -1.01523344 | Lipidmaps | M+H | C42H76NO8P | LMGP010116 47 | - | - | - | - |
| 130 | PC(18:3(9Z,12Z,15Z)/14:0) | PC | 728.52181 | 19.90475 | 2b | 19.90_728.5218m/z | 1598 | | -0.92293221 | Lipidmaps | M+H | C40H74NO8P | LMGP010116 73 | - | - | - | - |
| 131 | PC(18:3(9Z,12Z,15Z)/16:1(9Z)) | PC | 754.537086 | 20.65995 | 2b | 20.66_753.5298n | 1789 | 753.52981 | -1.3870781 | Lipidmaps | M+H, M+Na | C42H76NO8P | LMGP010116 78 | - | - | - | - |

| | | | | | | | | | | | | | | | | | |
|---|---|---|---|---|---|---|---|---|---|---|---|---|---|---|---|---|---|
| 132 | PC(18:3(9Z,12Z,15Z)/18:2(9Z,12Z)) | PC | 780.553924 | 20.5385 | 2b | 20.54_780.5539m/z | 182 | | 0.183121373 | Lipidmaps | M+H | C44H78NO8P | LMGP010116 83 | - | - | - | - |
| 133 | PC(16:0/18:1(9Z)) | PC | 760.584603 | 22.2077 | 2a | 22.21_760.5846m/z | 2166 | | -0.63018833 | Mona | M+H | C42H82NO8P | | - | - | - | - |
| 134 | PE 34:2 | PE | 716.522728 | 21.53395 | 2a | 21.53_715.5155n | 239 | 715.515451 | 0.344096319 | Metlin | M+H, M+Na | C39H74NO8P | | - | - | - | - |
| 135 | GPEtn(15:0/18:3) | PE | 700.491573 | 20.41041667 | 2b | 20.41_700.4916m/z | 1690 | | 0.560361628 | Lipidblast | M+H | C38H70NO8P | | - | - | - | - |
| 136 | GPEtn(18:2/15:0) | PE | 702.506537 | 21.047 | 2b | 21.05_702.5065m/z | 240 | | -0.41957423 | Lipidblast | M+H | C38H72NO8P | | - | - | - | - |
| 137 | GPEtn(18:2/18:0) | PE | 744.553129 | 22.4826 | 2b | 22.48_744.5531m/z | 2213 | | -0.87777274 | Lipidblast | M+H | C41H78NO8P | | - | - | - | - |
| 138 | GPEtn(18:2/18:1) | PE | 742.537196 | 21.80765 | 2b | 21.81_742.5372m/z | 2094 | | -1.26226628 | Lipidblast | M+H | C41H76NO8P | | - | - | - | - |
| 139 | GPEtn(18:2/18:2) | PE | 740.522953 | 21.13015 | 2b | 21.13_739.5157n | 1915 | 739.515677 | 0.638307439 | Lipidblast | M+H, M+Na | C41H74NO8P | | - | - | - | - |
| 140 | GPEtn(18:2/20:2) | PE | 768.551281 | 21.98491667 | 2b | 21.98_768.5513m/z | 2124 | | -3.25782016 | Lipidblast | M+H | C43H78NO8P | | - | - | - | - |
| 141 | GPEtn(18:3/16:0) | PE | 714.5072 | 20.92943333 | 2b | 20.93_714.5072m/z | 198 | | 0.516476217 | Lipidblast | M+H | C39H72NO8P | | - | - | - | - |
| 142 | GPEtn(18:3/18:2) | PE | 738.507125 | 20.5077 | 2b | 20.51_738.5071m/z | 1757 | | 0.398376851 | Lipidblast | M+H | C41H72NO8P | | - | - | - | - |
| 143 | GPEtn(18:3/18:3) | PE | 736.491444 | 19.89436667 | 2b | 19.89_736.4914m/z | 1595 | | 0.356601389 | Lipidblast | M+H | C41H70NO8P | | - | - | - | - |
| 144 | PG(20:3(8Z,11Z,14Z)/14:1(9Z)) | PG | 765.46735 | 20.09925 | 2b | 20.10_765.4673m/z | 1639 | | -0.47931704 | Lipidmaps | M+Na | C40H71O10P | LMGP040105 92 | - | - | - | - |
| 145 | PG(22:1(11Z)/20:5(5Z,8Z,11Z,14Z,17Z)) | PG | 873.553135 | 24.9434 | 2b | 24.94_873.5531m/z | 2590 | | -9.95944784 | Lipidmaps | M+Na | C48H83O10P | LMGP040107 49 | - | - | - | - |
| 146 | PG(22:2(13Z,16Z)/17:2(9Z,12Z)) | PG | 813.566899 | 25.83483333 | 2b | 25.83_813.5669m/z | 2693 | | 3.55261813 | Lipidmaps | M+H | C45H81O10P | LMGP040107 66 | - | - | - | - |
| 147 | PG(16:0/16:0) | PG | 745.499768 | 22.6832 | 2b | 22.68_745.4998m/z | 236 | | 1.054516366 | Lipidmaps | M+Na | C38H75O10P | LMGP040109 86 | - | - | - | - |
| 148 | PI(16:0/22:6(4Z,7Z,10Z,13Z,16Z,19Z)) | PI | 865.525483 | 24.61073333 | 2b | 24.61_865.5255m/z | 2558 | | 3.333773567 | Lipidmaps | M+H-H2O | C47H79O13P | LMGP060100 11 | - | - | - | - |
| 149 | PI(22:1(11Z)/18:4(6Z,9Z,12Z,15Z)) | PI | 935.569694 | 20.63283333 | 2b | 20.63_912.5804n | 1784 | 912.580428 | 8.381332622 | Lipidmaps | M+NH4, M+Na | C49H85O13P | LMGP060107 10 | - | - | - | - |
| 150 | PI(22:2(13Z,16Z)/16:1(9Z)) | PI | 871.573174 | 25.40473333 | 2b | 25.40_871.5732m/z | 2659 | | 4.14496528 | Lipidmaps | M+H-H2O | C47H85O13P | LMGP060107 32 | - | - | - | - |
| 151 | PI(22:4(7Z,10Z,13Z,16Z)/16:1(9Z)) | PI | 867.540078 | 24.85003333 | 2b | 24.85_867.5401m/z | 2578 | | 2.13313165 | Lipidmaps | M+H-H2O | C47H81O13P | LMGP060107 63 | - | - | - | - |
| 152 | Similar to PGF2alpha methyl ether | Prostaglandins | 337.273607 | 21.54025 | 2b | 21.54_337.2736m/z | 104 | | -0.32204854 | Lipidmaps | M+H-H2O | C21H38O4 | LMFA030100 73 | - | - | - | - |
| 153 | Similar to PGF2alpha methyl ether | Prostaglandins | 337.273591 | 22.34921667 | 2b | 22.35_337.2736m/z | 2189 | | -0.36748054 | Lipidmaps | M+H-H2O | C21H38O4 | LMFA030100 73 | - | - | - | - |
| 154 | PS(15:0/21:0) | PS | 774.562364 | 19.1027 | 2b | 19.10_774.5624m/z | 1472 | | -2.50401814 | Lipidmaps | M+H-H2O | C42H82NO10P | LMGP030109 11 | - | - | - | - |
| 155 | Chelidonic acid | Pyran | 185.007978 | 10.28493333 | 2a | 10.28_185.0080m/z | 783 | | -0.46965486 | Mona | M+H | C7H4O6 | | - | - | - | - |
| 156 | Muurolladie-3-One | Sesquiterpenoid | 219.174289 | 22.79603333 | 2a | 22.80_219.1743m/z | 2287 | | -0.24146933 | Mona | M+H | C15H22O | | - | - | - | - |
| 157 | Nootkatone | Sesquiterpenoid | 219.174398 | 13.73608333 | 2a | 13.74_219.1744m/z | 267 | | 0.256086158 | Mona | M+H | C15H22O | | - | - | - | - |
| 158 | Phytosphingosine | Sphingoid | 318.300317 | 9.654 | 2a | 9.65_318.3003m/z | 284 | | 0.147376885 | Mona | M+H | C18H39NO3 | | - | - | - | - |
| 159 | Similar to Stigmasterol | Sterol | 395.367487 | 22.49873333 | 2a | 22.50_395.3675m/z | 110 | | 0.628127717 | Metlin | M+H-H2O | C29H48O | | - | - | - | - |
| 160 | Similar to beta-Sitosterol | Sterol | 397.382783 | 18.93543333 | 2a | 18.94_397.3828m/z | 1450 | | -0.23041992 | Metlin | M+H-H2O | C29H50O | | - | - | - | - |
| 161 | Similar to beta-Sitosterol | Sterol | 397.383109 | 22.6832 | 2a | 22.68_397.3831m/z | 2255 | | 0.557653696 | Metlin | M+H-H2O | C29H50O | | - | - | - | - |
| 162 | beta-Sitostenone | Sterol | 413.377636 | 23.08555 | 2a | 23.09_413.3776m/z | 2357 | | -0.3787738 | Metlin | M+H | C29H48O | | - | - | - | - |
| 163 | Similar to Stigmasterol | Sterol | 395.367222 | 18.07453333 | 2a | 18.07_395.3672m/z | 1331 | | -0.01202439 | In-house | M+H-H2O | C29H48O | | - | - | - | - |
| 164 | Similar to Stigmasterol | Sterol | 395.367252 | 21.97848333 | 1a | 21.98_395.3673m/z | 2122 | | 0.057313143 | In-house | M+H-H2O | C29H48O | | - | - | - | - |
| 165 | Episterol | Sterol | 381.351479 | 21.35743333 | 2a | 21.36_381.3515m/z | 1986 | | -0.24670355 | In-house | M+H-H2O | C28H46O | | - | - | - | - |
| 166 | Similar to Pheophytin A | Tetrapyrroles and derivatives | 871.57336 | 25.48563333 | 2a | 25.49_871.5734m/z | 2664 | | 0.186367913 | Mona | M+H | C55H74N4O5 | | - | - | - | - |
| 167 | Similar to Pheophytin A | Tetrapyrroles and derivatives | 871.573463 | 25.60653333 | 2a | 25.61_870.5662n | 2679 | 870.566186 | 0.303890114 | Mona | M+H, M+NH4, M+Na | C55H74N4O5 | | - | - | - | - |
| 168 | 3-Hydroxyechinenone | Tetraterpenoids | 567.419599 | 19.8286 | 2a | 19.83_567.4196m/z | 1552 | | -0.10368103 | Mona | M+H | C40H54O2 | | - | - | - | - |
| 169 | Antheraxanthin | Tetraterpenoids | 585.430756 | 20.41673333 | 2a | 20.42_585.4308m/z | 1722 | | 0.912992007 | Mona | M+H | C40H56O3 | | - | - | - | - |
| 170 | Canthaxanthin (Euglenanone) | Tetraterpenoids | 565.40399 | 17.39915 | 2a | 17.40_564.3973n | 1239 | 564.397295 | 0.999507953 | Mona | M+H-H2O, M+H, M+NH4 | C40H52O2 | | - | - | - | - |
| 171 | Similar to Echinenone | TetraterpenoidsÂ | 568.427216 | 20.41041667 | 2a | 20.41_550.4179n | 1688 | 550.417867 | 0.727865033 | Mona | M+H-H2O, M+H, M+NH4 | C40H54O | | - | - | - | - |
| 172 | Similar to Echinenone | TetraterpenoidsÂ | 568.427316 | 21.06753333 | 2a | 21.07_550.4179n | 103 | 550.417867 | 0.728472706 | Mona | M+H-H2O, M+H, M+NH4 | C40H54O | | - | - | - | - |
| 173 | Phylloquinone | Vitamin | 451.35716 | 23.75835 | 2a | 23.76_451.3572m/z | 2465 | | 0.228100927 | Metlin | M+H | C31H46O2 | | - | - | - | - |
| 174 | Similar to Nostoxanthin | Xanthophyll | 601.42493 | 18.88435 | 2a | 18.88_600.4178n | 97 | 600.417847 | -0.02169119 | Mona | M+H-H2O, M+H | C40H56O4 | | - | - | - | - |
| 175 | Similar to Nostoxanthin | Xanthophyll | 601.425063 | 19.38785 | 2a | 19.39_601.4251m/z | 50 | | -0.1223843 | Mona | M+H | C40H56O4 | | - | - | - | - |
| 176 | NCGC00385443-01!(2S)-3-(3,4-dihydroxyphenyl)-2-[[(E)-3-(3,4-dihydroxyphenyl)prop-2-enoyl]amino]propanoic acid | | 360.10795 | 6.2053 | 2a | 6.21_360.1079m/z | 79 | | 0.477421972 | Mona | M+H | C18H17NO7 | | - | - | - | - |
| 177 | 2-(3_4-dihydroxyphenyl)-5_7-dihydroxy-6-[3_4_5-trihydroxy-6-(hydroxymethyl)oxan-2-yl]-4H-chromen-4-one | | 449.107264 | 5.751116667 | 1a | 5.75_449.1073m/z | 550 | | -2.25895966 | In-house | M+H | C21H20O11 | | - | - | - | - |
