## Supplementary material for "Untargeted metabolomics reveals key metabolites and genes underlying salinity tolerance mechanisms in maize": List of identified mass features and differentially abundant metabolites in roots of NC326 and C68 inbred lines under control and salinity conditions.

**Supplementary Dataset S2**: List of annotated mass features in roots in NC326 and C68 inbred lines and differentially abundant (DA) metabolites under control and salinity conditions

| Sr No. | Final Description | Class | m/z | Retention time (min) | Annotation level | Compound | ClusterID | Neutral mass (Da) | Mass Error (ppm) | MS2Match | Adducts | Formula | LipidMAPS | DA in Control_C68 vs NC326 | DA in Salinity_C68 vs NC326 | DA in C68_Control vs Salinity | DA in NC326_Control vs Salinity |
|---|---|---|---|---|---|---|---|---|---|---|---|---|---|---|---|---|---|
| 1 | LPC 18:3 | Lyso PC | 518.3240843 | 12.17655 | 2b | 12.18_518.3241m/z | 975 | | -0.061097486 | Lipidmaps | M+H | C26H48NO7P | LMGP01050128 | - | - | X | - |
| 2 | LPC 18:1 | Lyso PC | 522.35565 | 13.6902 | 2b | 13.69_522.3556m/z | 1036 | | 0.448606718 | Lipidmaps | M+H | C26H52NO7P | LMGP01050029 | - | - | X | X |
| 3 | LPC 18:2 | Lyso PC | 520.3403035 | 12.87763333 | 2b | 12.88_519.3330n | 17 | 519.3330271 | 1.035018843 | Lipidblast | M+H-H2O, M+H, M+Na | C26H50NO7P | | - | - | X | X |
| 4 | LPC 18:2 | Lyso PC | 520.340175 | 13.13598333 | 2b | 13.14_519.3329n | 30 | 519.3328986 | 0.787524254 | Lipidblast | M+H-H2O, M+H, M+Na | C26H50NO7P | | - | - | X | X |
| 5 | Similar to Feruloyl putrescine | Hydroxycinnamic acid | 265.1549534 | 3.13735 | 2a | 3.14_265.1550m/z | 381 | | 1.07676334 | Mona | M+H | C14H20N2O3 | | - | X | - | - |
| 6 | Similar to Feruloyl putrescine | Hydroxycinnamic acid | 265.1549321 | 3.39225 | 2a | 3.39_265.1549m/z | 396 | | 0.996197587 | Mona | M+H | C14H20N2O3 | | - | X | - | - |
| 7 | GPEtn(15:0/18:2) | PE | 702.5065781 | 21.03396667 | 2b | 21.03_701.4993n | 1620 | 701.4993017 | -0.360926472 | Lipidblast | M+H, M+Na | C38H72NO8P | | - | X | - | - |
| 8 | PE-NMe2(18:1(9Z)/16:0) | PE | 746.5668445 | 21.74816667 | 2b | 21.75_746.5668m/z | 1805 | | -3.470026053 | Lipidmaps | M+H | C41H80NO8P | LMGP02010330 | - | X | - | - |
| 9 | Pinolenic Acid methyl ester | FAME | 293.2477233 | 18.79 | 2a | 18.79_293.2477m/z | 1332 | | 0.741348898 | Metlin | M+H | C19H32O2 | | - | X | X | - |
| 10 | PG(18:2(9Z,12Z)/16:0) | PG | 769.4985116 | 23.16851667 | 2b | 23.17_769.4985m/z | 2194 | | -0.662165156 | Lipidmaps | M+Na | C40H75O10P | LMGP04010877 | - | X | X | - |
| 11 | GPEtn(16:0/18:3) | PE | 714.5069142 | 20.91931667 | 2b | 20.92_713.4996n | 1599 | 713.4996377 | 0.11610745 | Lipidblast | M+H, M+Na | C39H72NO8P | | X | - | - | - |
| 12 | GPEtn(18:2/18:0) | PE | 744.5534357 | 22.47555 | 2b | 22.48_744.5534m/z | 2009 | | -0.46512181 | Lipidblast | M+H | C41H78NO8P | | X | - | - | - |
| 13 | 5-Deoxy-5-(methylthio)adenosine | Purine | 298.0971209 | 3.7816 | 1a | 3.78_298.0971m/z | 243 | | 0.822929306 | in-house | M+H | C11H15N5O3S | | X | - | - | - |
| 14 | Coumaroyl tyramine | Hydroxycinnamic acid | 284.1283436 | 7.871966667 | 2a | 7.87_283.1211n | 213 | 283.1210672 | 0.790455755 | Mona | M+H, M+Na | C17H17NO3 | | X | - | - | X |
| 15 | Glutathione reduced | Antioxidant | 308.091502 | 1.649966667 | 2a | 1.65_308.0915m/z | 255 | | 1.365338544 | Mona | M+H | C10H17N3O6S | | X | - | X | - |
| 16 | Moupinamide | Phenols | 314.1389906 | 8.015616667 | 1a | 8.02_314.1390m/z | 806 | | 2.280711864 | in-house | M+H | C18H19NO4 | | X | - | X | - |
| 17 | GDIMBOA | Carbohydrates conjugates?? | 366.0798739 | 3.971283333 | 2a | 3.97_366.0799m/z | 434 | | 0.938736123 | Mona | M+Na | C14H17NO9 | | X | X | - | - |
| 18 | DGDG(18:0/18:2) | DGDG | 967.6322901 | 22.45421667 | 2b | 22.45_967.6323m/z | 150 | | -0.585221914 | Lipidblast | M+Na | C51H92O15 | | X | X | - | - |
| 19 | 28:6(10Z,13Z,16Z,19Z,22Z,25Z) | Fatty acid | 413.345185 | 16.74838333 | 2b | 16.75_413.3452m/z | 184 | | 9.162159275 | Lipidmaps | M+H | C28H44O2 | LMFA01030839 | X | X | - | - |
| 20 | Similar to Feruloyl allylamine | Ferulic acid and derivatives | 234.112754 | 3.169583333 | 2a | 3.17_234.1128m/z | 387 | | 1.219330313 | Mona | M+H | C13H15NO3 | | X | X | - | - |
| 21 | Similar to Feruloyl allylamine | Ferulic acid and derivatives | 234.1127975 | 3.43185 | 2a | 3.43_234.1128m/z | 400 | | 1.405927486 | Mona | M+H | C13H15NO3 | | X | X | - | - |
| 22 | PC(18:2(9Z,12Z)/18:1(9Z)) | PC | 784.5848315 | 21.8037 | 2b | 21.80_783.5776n | 1815 | 783.5775551 | -0.319200109 | Lipidmaps | M+H, M+Na | C44H82NO8P | LMGP01011624 | X | X | - | - |
| 23 | (9R,13R)-10,11-dihydro-12-oxo-15-phytoenoic acid | 12-oxophytodienoic acid metabolites | 295.2266527 | 23.11143333 | 2b | 23.11_294.2195n | 2183 | 294.2194598 | -0.118888434 | Lipidmaps | M+H-H2O, M+H | C18H30O3 | LMFA02010006 | - | - | - | - |
| 24 | Similar to PGF2alpha methyl ether | Aliphatic alcohol | 337.2736287 | 16.29013333 | 2b | 16.29_354.2769n | 44 | 354.2769169 | -0.261908607 | Lipidmaps | M+H-H2O, M+H, M+Na | C21H38O4 | LMFA03010073 | - | - | - | - |
| 25 | Tetradecylamine | Amines | 214.2532288 | 8.762983333 | 2a | 8.76_214.2532m/z | 833 | | 1.417862656 | Metlin | M+H | C14H31N | | - | - | - | - |
| 26 | 2,2'-(Tetradecylimino)diethanol | Amino alcohol | 302.3055474 | 9.507483333 | 2a | 9.51_302.3055m/z | 880 | | 0.635465049 | Mona | M+H | C18H39NO2 | | - | - | - | - |
| 27 | N,N-dimethyl-Safingol | Amino alcohol | 330.3366873 | 11.35948333 | 2b | 11.36_329.3295n | 960 | 329.3295332 | 0.466316018 | Lipidmaps | M+H-H2O, M+H | C20H43NO2 | LMSP01080056 | - | - | - | - |
| 28 | Cer(t18:0/16:0) | Cer | 556.5278891 | 21.73698333 | 2b | 21.74_555.5206n | 1803 | 555.5206127 | -3.684790586 | Lipidmaps | M+H, M+Na | C34H69NO4 | LMSP02030001 | - | - | - | - |
| 29 | Trans-Ferulic acid | Cinnamic acids and derivatives | 177.0557162 | 24.89493333 | 2a | 24.89_177.0557m/z | 2387 | | 5.645742417 | in-house | M+H-H2O | C10H10O4 | | - | - | - | - |
| 30 | Coumarin | Coumarins and derivatives | 147.0446012 | 7.87635 | 2a | 7.88_147.0446m/z | 213 | | 3.734275916 | Metlin | M+H | C9H6O2 | | - | - | - | - |
| 31 | 1-methyl-cyclopentanol | Cyclopentanols | 118.1231906 | 23.91478333 | 2b | 23.91_118.1232m/z | 2315 | | 5.495413182 | Lipidmaps | M+NH4 | C6H12O | LMFA05000538 | - | - | - | - |
| 32 | DG(17:1(9Z)/17:2(9Z,12Z)/0:0)[iso2] | DG | 613.4799547 | 23.83675 | 2b | 23.84_613.4800m/z | 61 | | -0.493223959 | Lipidmaps | M+Na | C37H66O5 | LMGL02010024 | - | - | - | - |
| 33 | Similar to DG(16:0/18:2(9Z,12Z)/0:0)[iso2] | DG | 575.5031468 | 22.70735 | 2b | 22.71_592.5064n | 2113 | 592.506435 | -0.405472942 | Lipidmaps | M+H-H2O, M+H | C37H68O5 | LMGL02010027 | - | - | - | - |
| 34 | Similar to DG(16:0/18:2(9Z,12Z)/0:0)[iso2] | DG | 615.4952851 | 24.011 | 2b | 24.01_615.4953m/z | 158 | | -1.031049925 | Lipidmaps | M+Na | C37H68O5 | LMGL02010027 | - | - | - | - |
| 35 | DG(16:0/18:3(9Z,12Z,15Z)/0:0)[iso2] | DG | 591.4978556 | 20.91931667 | 2b | 20.92_591.4979m/z | 1598 | | -0.755435515 | Lipidmaps | M+H | C37H66O5 | LMGL02010032 | - | - | - | - |

| | | | | | | | | | | | | | | | | | |
|---|---|---|---|---|---|---|---|---|---|---|---|---|---|---|---|---|---|
| 36 | DG(18:1(9Z)/18:3(9Z,12Z,15Z)/0:0)[iso2] | DG | 639.4954128 | 23.70435 | 2b | 23.70_616.5064n | 2290 | 616.5064346 | -0.390398621 | Lipidmaps | M+H, M+NH4, M+Na | C39H68O5 | LMGL02010064 | - | - | - | - |
| 37 | DG(18:3(9Z,12Z,15Z)/18:3(9Z,12Z,15Z)/0:0) | DG | 595.469618 | 23.99971667 | 2b | 24.00_595.4696m/z | 109 | | -4.031002222 | Lipidmaps | M+H-H2O | C39H64O5 | LMGL02010079 | - | - | - | - |
| 38 | DG(15:1(9Z)/20:5(5Z,8Z,11Z,14Z,17Z)/0:0)[iso2] | DG | 581.4539537 | 23.32025 | 2b | 23.32_581.4540m/z | 110 | | -4.149219594 | Lipidmaps | M+H-H2O | C38H62O5 | LMGL02010465 | - | - | - | - |
| 39 | DG(18:4(6Z,9Z,12Z,15Z)/16:0/0:0)[iso2] | DG | 611.463911 | 23.36731667 | 2b | 23.37_611.4639m/z | 2240 | | -1.163705532 | Lipidmaps | M+Na | C37H64O5 | LMGL02010498 | - | - | - | - |
| 40 | DGDG(16:0/18:1) | DGDG | 941.6163047 | 22.1569 | 2b | 22.16_918.6271n | 1900 | 918.6271024 | -0.946841768 | Lipidblast | M+NH4, M+Na | C49H90O15 | | - | - | - | - |
| 41 | DGDG(16:0/18:2) | DGDG | 939.6007887 | 21.50598333 | 2b | 21.51_939.6008m/z | 105 | | -0.822737154 | Lipidblast | M+Na | C49H88O15 | | - | - | - | - |
| 42 | DGDG(18:1/18:2) | DGDG | 965.6161007 | 21.79728333 | 2b | 21.80_942.6269n | 1818 | 942.6268902 | -1.14783669 | Lipidblast | M+NH4, M+Na | C51H90O15 | | - | - | - | - |
| 43 | DGDG(18:2/20:0) | DGDG | 995.6629833 | 23.23495 | 2b | 23.23_972.6743n | 2206 | 972.6743274 | -0.611682265 | Lipidblast | M+NH4, M+Na | C53H96O15 | | - | - | - | - |
| 44 | DGTS(16:0/18:2(9Z,12Z)) | DGTS | 736.6073359 | 21.6826 | 2b | 21.68_736.6073m/z | 1783 | | -1.691725577 | Lipidmaps | M+H | C44H81NO7 | LMGL00000121 | - | - | - | - |
| 45 | 8,15-Pimaradiene | Diterpenoids | 273.2575777 | 17.66996667 | 2a | 17.67_273.2576m/z | 1243 | | -0.36644267 | Metlin | M+H | C20H32 | | - | - | - | - |
| 46 | Similar to Linolenic acid | Fatty acid | 279.2322485 | 14.04515 | 2a | 14.05_278.2247n | 16 | 278.2246657 | 0.307199888 | Mona | M+H-H2O, M+H | C18H30O2 | | - | - | - | - |
| 47 | 26:2(5Z,9Z)(25Me) | Fatty acid | 429.3699884 | 23.04363333 | 2b | 23.04_429.3700m/z | 2169 | | -0.770493703 | Lipidmaps | M+Na | C27H50O2 | LMFA01020359 | - | - | - | - |
| 48 | 13Z,17-Octadecadiene-9,11-diynoic acid | Fatty acid | 295.1655807 | 1.698233333 | 2b | 1.70_295.1656m/z | 317 | | -4.665879553 | Lipidmaps | M+Na | C18H24O2 | LMFA01030510 | - | - | - | - |
| 49 | 5Z,8Z,14Z-Eicosatrien-11-ynoic acid | Fatty acid | 285.2215993 | 17.18218333 | 2b | 17.18_302.2249n | 1225 | 302.2248876 | 1.016997417 | Lipidmaps | M+H-H2O, M+H | C20H30O2 | LMFA01030693 | - | - | - | - |
| 50 | beta-kamlolenic acid | Fatty acid | 295.2265584 | 22.5873 | 2b | 22.59_294.2196n | 111 | 294.2196486 | 0.522673354 | Lipidmaps | M+H-H2O, M+H | C18H30O3 | LMFA02000157 | - | - | - | - |
| 51 | 10(E)_12(Z)-Conjugated Linoleic Acid | Fatty acid | 263.2374501 | 17.69111667 | 1a | 17.69_280.2407n | 1 | 280.2407384 | 1.921169925 | in-house | M+H-H2O, M+H | C18H32O2 | | - | - | - | - |
| 52 | Similar to Linolenic acid | Fatty acid | 279.2321333 | 15.21281667 | 2a | 15.21_279.2321m/z | 1113 | | 0.923415113 | in-house | M+H | C18H30O2 | | - | - | - | - |
| 53 | Linoleoyl Ethanolamide | Fatty acid amide | 324.2901007 | 15.1734 | 1a | 15.17_324.2901m/z | 1112 | | 1.312525115 | in-house | M+H | C20H37NO2 | | - | - | - | - |
| 54 | Oleoyl Ethanolamide | Fatty acid amide | 326.3058058 | 16.11825 | 1a | 16.12_326.3058m/z | 1161 | | 1.319999439 | in-house | M+H | C20H39NO2 | | - | - | - | - |
| 55 | Methyl linoleate | Fatty acid ester | 295.2633288 | 19.57896667 | 2a | 19.58_295.2633m/z | 68 | | 0.584561322 | Metlin | M+H | C19H34O2 | | - | - | - | - |
| 56 | (E)-9-Octadecenoic acid methyl ester | Fatty acid ester | 297.2790555 | 20.46083333 | 2a | 20.46_297.2791m/z | 167 | | 0.839180901 | Metlin | M+H | C19H36O2 | | - | - | - | - |
| 57 | Juniperic acid | Fatty Acyls | 290.2692872 | 8.613833333 | 2b | 8.61_290.2693m/z | 828 | | 1.163607034 | Lipidmaps | M+NH4 | C16H32O3 | LMFA01050051 | - | - | - | - |
| 58 | 10-oxo-nonadecanoic acid | Fatty Acyls | 335.2581954 | 21.23863333 | 2b | 21.24_335.2582m/z | 1678 | | 8.101138621 | Lipidmaps | M+Na | C19H36O3 | LMFA01060128 | - | - | - | - |
| 59 | gamma- 12,13-DiHODE | Fatty Acyls | 295.2270006 | 14.6375 | 2b | 14.64_295.2270m/z | 73 | | 0.734678634 | Lipidmaps | M+H-H2O | C18H32O4 | LMFA02000050 | - | - | - | - |
| 60 | 12-OPDA | Fatty Acyls?? | 293.2113005 | 13.45546667 | 2a | 13.46_293.2113m/z | 1032 | | 0.613809613 | Metlin | M+H | C18H28O3 | | - | - | - | - |
| 61 | 6,8,10,12-pentadecatetraenal | Fatty aldehyde | 219.1745033 | 22.77843333 | 2b | 22.78_219.1745m/z | 2124 | | 0.740347557 | Lipidmaps | M+H | C15H22O | LMFA06000087 | - | - | - | - |
| 62 | Glucocerebroside | Glucocerebroside | 696.5409802 | 20.02251667 | 2a | 20.02_713.5443n | 26 | 713.5442684 | 0.11975858 | Metlin | M+H-H2O, M+H, M+Na | C40H75NO9 | | - | - | - | - |
| 63 | 2-Linoleoyl Glycerol | Glycerin fatty acid ester | 355.2831709 | 16.1161 | 1a | 16.12_354.2770n | 45 | 354.2770124 | 0.034996054 | in-house | M+H-H2O, M+H | C21H38O4 | | - | - | - | - |
| 64 | Similar to 2-Linoleoyl Glycerol | Glycerin fatty acid ester | 337.2739246 | 21.53193333 | 2a | 21.53_337.2739m/z | 59 | | 0.600788127 | in-house | M+H-H2O | C21H38O4 | | - | - | - | - |
| 65 | Similar to 2-Linoleoyl Glycerol | Glycerin fatty acid ester | 337.2736922 | 22.346 | 2a | 22.35_337.2737m/z | 1967 | | -0.055218387 | in-house | M+H-H2O | C21H38O4 | | - | - | - | - |
| 66 | Similar to 2-Linoleoyl Glycerol | Glycerin fatty acid ester | 337.273886 | 22.70735 | 2a | 22.71_337.2739m/z | 100 | | 0.491764408 | in-house | M+H-H2O | C21H38O4 | | - | - | - | - |
| 67 | 1_2-Dioleoyl-sn-glycerol | Glycerolipids | 603.5334062 | 22.47296667 | 2a | 22.47_603.5334m/z | 1989 | | -2.103844234 | in-house | M+H-H2O | C39H72O5 | | - | - | - | - |
| 68 | Monooleoylglycerol | Glycerolipids | 339.2910447 | 22.1569 | 2a | 22.16_339.2910m/z | 1897 | | 3.741116428 | in-house | M+H-H2O | C21H40O4 | | - | - | - | - |
| 69 | 9Z-Nonadecene | Hydrocarbon | 284.3313292 | 11.29438333 | 2b | 11.29_284.3313m/z | 957 | | 0.572502945 | Lipidmaps | M+NH4 | C19H38 | LMFA11000112 | - | - | - | - |
| 70 | 3-Indoleacrylic acid | Indoles | 188.0711635 | 3.7328 | 2a | 3.73_187.0639n | 89 | 187.0638871 | 2.98574611 | Mona | M+H, M+NH4 | C11H9NO2 | | - | - | - | - |
| 71 | LPC 16:0 | Lyso PC | 496.3401063 | 13.41828333 | 2a | 13.42_495.3328n | 200 | 495.3328299 | 0.687082866 | Mona | M+H-H2O, M+H, M+Na | C24H50NO7P | | - | - | - | - |
| 72 | LPE 16:0 | Lyso PE | 454.2928716 | 13.35283333 | 2a | 13.35_453.2855n | 1030 | 453.2855035 | -0.079100537 | Metlin | M+H-H2O, M+H, M+Na | C21H44NO7P | | - | - | - | - |
| 73 | LPE 18:2 | Lyso PE | 478.2933508 | 12.80875 | 2a | 12.81_477.2835n | 29 | 477.2835357 | -4.198159827 | Mona | M+H-H2O, M+H, M+Na | C23H44NO7P | | - | - | - | - |
| 74 | PI(18:2(9Z,12Z)/0:0) | Lyso PI | 619.285358 | 18.05681667 | 2b | 18.06_596.2966n | 180 | 596.2965944 | 0.722369395 | Lipidmaps | M+H-H2O, M+H, M+Na | C27H49O12P | LMGP06050010 | - | - | - | - |
| 75 | Similar to S-cucujolide III | Macrolide | 247.1667892 | 21.93003333 | 2b | 21.93_247.1668m/z | 1852 | | -0.274485907 | Lipidmaps | M+Na | C14H24O2 | LMFA07040046 | - | - | - | - |

| | | | | | | | | | | | | | | | | | |
|---|---|---|---|---|---|---|---|---|---|---|---|---|---|---|---|---|---|
| 76 | Similar to S-cucujolide III | Macrolide | 247.1667648 | 22.80861667 | 2b | 22.81_247.1668m/z | 2128 | | -0.383170952 | Lipidmaps | M+Na | C14H24O2 | LMFA07040046 | - | - | - | - |
| 77 | Similar to S-cucujolide III | Macrolide | 247.1667154 | 23.3504 | 2b | 23.35_247.1667m/z | 2232 | | -0.603595254 | Lipidmaps | M+Na | C14H24O2 | LMFA07040046 | - | - | - | - |
| 78 | MGDG(18:0(9Z)/18:2(9Z,12Z)) | MGDG | 805.5789313 | 23.4343 | 2b | 23.43_782.5898n | 2252 | 782.5897889 | -1.290493847 | Lipidmaps | M+NH4, M+Na | C45H82O10 | LMGL05010022 | - | - | - | - |
| 79 | MGDG(16:0/18:2(9Z,12Z)) | MGDG | 777.5487343 | 22.70006667 | 2b | 22.70_777.5487m/z | 57 | | 0.019715994 | Lipidmaps | M+Na | C43H78O10 | LMGL05010026 | - | - | - | - |
| 80 | MGDG(18:1/16:0) | MGDG | 779.5638024 | 23.24425 | 2b | 23.24_779.5638m/z | 2211 | | -0.749439895 | Lipidblast | M+Na | C43H80O10 | | - | - | - | - |
| 81 | MGDG(18:1/18:2) | MGDG | 803.5637292 | 22.91621667 | 2b | 22.92_803.5637m/z | 2147 | | -0.820201798 | Lipidblast | M+Na | C45H80O10 | | - | - | - | - |
| 82 | N-Phenyl-1-naphthylamine | Naphthalenes | 220.1123298 | 14.7173 | 2a | 14.72_220.1123m/z | 1083 | | 1.159095164 | Mona | M+H | C16H13N | | - | - | - | - |
| 83 | 1-(2-Pyrimidyl)piperazine | N-arylpiperazine | 165.1132802 | 19.92586667 | 2a | 19.93_165.1133m/z | 1433 | | -1.173702772 | Metlin | M+H | C8H12N4 | | - | - | - | - |
| 84 | Triphenyl phosphate | Organophosphates | 327.0783977 | 13.94161667 | 2a | 13.94_327.0784m/z | 1047 | | 0.998768381 | Metlin | M+H | C18H15O4P | | - | - | - | - |
| 85 | 13-HOTrE-LIKE | Oxylipin | 277.2161261 | 14.50615 | 2a | 14.51_294.2197n | 72 | 294.2197012 | 0.701506607 | Metlin | M+H-H2O, M+H, M+Na | C18H30O3 | | - | - | - | - |
| 86 | 13S-HpOTrE(gamma) | Oxylipin | 293.2113158 | 13.55733333 | 2b | 13.56_310.2146n | 1034 | 310.2146041 | 0.627462442 | Lipidmaps | M+H-H2O, M+Na | C18H30O4 | LMFA02000112 | - | - | - | - |
| 87 | PC(18:3(6Z,9Z,12Z)/18:3(6Z,9Z,12Z)) | PC | 778.5376191 | 19.90735 | 2a | 19.91_777.5303n | 1431 | 777.5303427 | -0.658944004 | Metlin | M+H, M+Na | C44H76NO8P | | - | - | - | - |
| 88 | GPCho(18:2/16:0) | PC | 780.5513236 | 21.58688333 | 2b | 21.59_780.5513m/z | 1764 | | -0.068953342 | Lipidblast | M+Na | C42H80NO8P | | - | - | - | - |
| 89 | GPCho(18:2/18:0) | PC | 808.5826511 | 22.48068333 | 2b | 22.48_808.5827m/z | 2016 | | -0.031603045 | Lipidblast | M+Na | C44H84NO8P | | - | - | - | - |
| 90 | GPCho(18:2/18:3) | PC | 802.5353956 | 20.55796667 | 2b | 20.56_802.5354m/z | 1540 | | -0.423535712 | Lipidblast | M+Na | C44H78NO8P | | - | - | - | - |
| 91 | PC(14:0/18:2(11Z,14Z)) | PC | 730.5384215 | 20.55796667 | 2b | 20.56_730.5384m/z | 1537 | | 0.397598837 | Lipidmaps | M+H | C40H76NO8P | LMGP01010494 | - | - | - | - |
| 92 | PC(16:0/18:2(11Z,13Z)) | PC | 758.5696352 | 21.54523333 | 2b | 21.55_758.5696m/z | 1754 | | 0.268835119 | Lipidmaps | M+H | C42H80NO8P | LMGP01010586 | - | - | - | - |
| 93 | PC(18:1(9Z)/18:3(9Z,12Z,15Z)) | PC | 782.5692801 | 21.1569 | 2b | 21.16_781.5620n | 1651 | 781.5620036 | -0.193871236 | Lipidmaps | M+H, M+Na | C44H80NO8P | LMGP01010898 | - | - | - | - |
| 94 | PC(18:3(9Z,12Z,15Z)/15:0) | PC | 742.5368124 | 20.56266667 | 2b | 20.56_741.5295n | 1538 | 741.5295359 | -1.778875754 | Lipidmaps | M+H, M+Na | C41H76NO8P | LMGP01011675 | - | - | - | - |
| 95 | PC(18:3(9Z,12Z,15Z)/16:0) | PC | 756.5538454 | 20.93913333 | 2b | 20.94_755.5466n | 170 | 755.546569 | 0.08459172 | Lipidmaps | M+H, M+Na | C42H78NO8P | LMGP01011677 | - | - | - | - |
| 96 | PC(18:3(9Z,12Z,15Z)/17:0) | PC | 770.5686752 | 21.43508333 | 2b | 21.44_770.5687m/z | 1729 | | -0.982865276 | Lipidmaps | M+H | C43H80NO8P | LMGP01011679 | - | - | - | - |
| 97 | PC(P-18:0/20:2(11Z,14Z)) | PC | 820.6268111 | 22.95268333 | 2b | 22.95_797.6374n | 162 | 797.6374066 | 9.485358218 | Lipidmaps | M+H-H2O, M+H, M+Na | C46H88NO7P | LMGP01030066 | - | - | - | - |
| 98 | PC(16:0/20:4(6E,8Z,11Z,14Z)(5OH[S])) | PC | 798.5637795 | 18.61706667 | 2b | 18.62_797.5560n | 122 | 797.5559516 | -1.401968125 | Lipidmaps | M+H-H2O, M+H | C44H80NO9P | LMGP20010003 | - | - | - | - |
| 99 | PC(16:0/9:0(CHO)) | PC | 650.4389181 | 15.90676667 | 2b | 15.91_649.4316n | 1145 | 649.4316417 | -0.35047285 | Lipidmaps | M+H, M+Na | C33H64NO9P | LMGP20010008 | - | - | - | - |
| 100 | PC(16:0/18:1(9Z)) | PC | 760.5846688 | 22.21108333 | 2a | 22.21_760.5847m/z | 1917 | | -0.543483733 | Mona | M+H | C42H82NO8P | | - | - | - | - |
| 101 | PC 16:0_16:0 | PC | 734.5677528 | 21.91325 | 1a | 21.91_734.5678m/z | 1850 | | -2.349669244 | in-house | M+H | C40H80NO8P | | - | - | - | - |
| 102 | PAz-PC | PC | 666.4340493 | 15.04516667 | 1a | 15.05_665.4268n | 191 | 665.4267729 | -0.04070012 | in-house | M+H, M+Na | C33H64NO10P | | - | - | - | - |
| 103 | GPEtn(16:1/16:0) | PE | 690.506347 | 21.24896667 | 2b | 21.25_690.5063m/z | 1685 | | -0.702434049 | Lipidblast | M+H | C37H72NO8P | | - | - | - | - |
| 104 | GPEtn(16:1/16:1) | PE | 688.4918677 | 20.53743333 | 2b | 20.54_687.4846n | 1531 | 687.4845913 | 0.998465863 | Lipidblast | M+H, M+Na | C37H70NO8P | | - | - | - | - |
| 105 | GPEtn(17:1/17:1) | PE | 716.5229807 | 21.52488333 | 2b | 21.52_716.5230m/z | 1747 | | 0.697885937 | Lipidblast | M+H | C39H74NO8P | | - | - | - | - |
| 106 | GPEtn(18:2/17:0) | PE | 730.5374672 | 21.99371667 | 2b | 21.99_730.5375m/z | 1867 | | -0.910498363 | Lipidblast | M+H | C40H76NO8P | | - | - | - | - |
| 107 | GPEtn(18:2/18:3) | PE | 738.5099753 | 20.51153333 | 2b | 20.51_737.5027n | 1524 | 737.5026989 | 4.262995656 | Lipidblast | M+H, M+Na | C41H72NO8P | | - | - | - | - |
| 108 | GPEtn(18:2/20:2) | PE | 768.5528705 | 21.97138333 | 2b | 21.97_767.5456n | 1861 | 767.545594 | -1.1869634 | Lipidblast | M+H, M+Na | C43H78NO8P | | - | - | - | - |
| 109 | GPEtn(18:3/16:0) | PE | 714.50648 | 18.99725 | 2b | 19.00_714.5065m/z | 65 | | -0.492344562 | Lipidblast | M+H | C39H72NO8P | | - | - | - | - |
| 110 | PE(16:0/18:2(9Z,12Z)) | PE | 738.5045922 | 21.5162 | 2b | 21.52_738.5046m/z | 1749 | | 0.232744064 | Lipidmaps | M+Na | C39H74NO8P | LMGP02010042 | - | - | - | - |
| 111 | PE(16:0/20:4(5Z,8Z,11Z,14Z)) | PE | 740.5228776 | 21.13668333 | 2b | 21.14_739.5156n | 1643 | 739.5156012 | 0.535748594 | Lipidmaps | M+H, M+Na | C41H74NO8P | LMGP02010096 | - | - | - | - |
| 112 | PE(18:2(9Z,12Z)/17:0) | PE | 730.5373117 | 21.70928333 | 2b | 21.71_730.5373m/z | 1794 | | -1.123739429 | Lipidmaps | M+H | C40H76NO8P | LMGP02010660 | - | - | - | - |
| 113 | PE(20:0/20:3(8Z,11Z,14Z)) | PE | 820.5903387 | 22.48583333 | 2b | 22.49_820.5903m/z | 2029 | | 9.607380104 | Lipidmaps | M+Na | C45H84NO8P | LMGP02010835 | - | - | - | - |
| 114 | PE(22:1(11Z)/20:0) | PE | 852.6525434 | 22.86633333 | 2b | 22.87_829.6634n | 99 | 829.6634117 | 8.866573211 | Lipidmaps | M+H, M+Na | C47H92NO8P | LMGP02011059 | - | - | - | - |
| 115 | Triphenylphosphine oxide | Phosphine oxide?? | 279.0936399 | 22.65946667 | 2a | 22.66_279.0936m/z | 2076 | | 1.121104914 | Mona | M+H | C18H15OP | | - | - | - | - |

| | | | | | | | | | | | | | | | | | |
|---|---|---|---|---|---|---|---|---|---|---|---|---|---|---|---|---|---|
| 116 | PI(22:2(13Z,16Z)/16:1(9Z)) | PI | 871.572352 | 25.44035 | 2b | 25.44_871.5724m/z | 2398 | | 3.219588402 | Lipidmaps | M+H-H2O | C47H85O13P | LMGP06010732 | - | - | - | - |
| 117 | PI(P-16:0/19:0) | PI | 854.6111481 | 22.47296667 | 2b | 22.47_854.6111m/z | 1991 | | -0.648132561 | Lipidmaps | M+NH4 | C44H85O12P | LMGP06030017 | - | - | - | - |
| 118 | Similar to Farnesyl acetone | Prenol lipids | 263.2371646 | 16.29013333 | 2a | 16.29_262.2303n | 44 | 262.2302925 | 2.39078128 | Metlin | M+H-H2O, M+H | C18H30O | | - | - | - | - |
| 119 | Similar to Farnesyl acetone | Prenol lipids | 263.2371363 | 19.58913333 | 2a | 19.59_262.2299n | 68 | 262.2299109 | 0.935533284 | Metlin | M+H-H2O, M+H | C18H30O | | - | - | - | - |
| 120 | PS(O-18:0/13:0) | PS | 708.511248 | 24.21768333 | 2b | 24.22_708.5112m/z | 2355 | | -8.689691042 | Lipidmaps | M+H | C37H74NO9P | LMGP03020019 | - | - | - | - |
| 121 | PS(O-20:0/20:3(8Z,11Z,14Z)) | PS | 810.6049999 | 21.46985 | 2b | 21.47_810.6050m/z | 1740 | | 5.157250375 | Lipidmaps | M+H-H2O | C46H86NO9P | LMGP03020062 | - | - | - | - |
| 122 | PS(P-20:0/22:2(13Z,16Z)) | PS | 838.6375685 | 22.47555 | 2b | 22.48_838.6376m/z | 2018 | | 6.470763912 | Lipidmaps | M+H-H2O | C48H90NO9P | LMGP03030084 | - | - | - | - |
| 123 | Chelidonic acid | Pyrans | 185.0085076 | 10.1347 | 2a | 10.13_185.0085m/z | 47 | | 2.409230736 | Mona | M+H | C7H4O6 | | - | - | - | - |
| 124 | Similar to Fusaric acid | Pyridines | 180.102543 | 5.839533333 | 2a | 5.84_179.0954n | 658 | 179.0953817 | 4.204711495 | Mona | M+H-H2O, M+H | C10H13NO2 | | - | - | - | - |
| 125 | Similar to Fusaric acid | Pyridines | 180.1024688 | 7.3256 | 2a | 7.33_179.0952n | 784 | 179.0951725 | 3.036563847 | Mona | M+H-H2O, M+H | C10H13NO2 | | - | - | - | - |
| 126 | Phytosphingosine | Sphingoid | 318.3007118 | 9.635683333 | 2a | 9.64_317.2929n | 209 | 317.2928541 | -0.44126195 | Mona | M+H-H2O, M+H | C18H39NO3 | | - | - | - | - |
| 127 | SPHINGANINE | Sphingoid | 302.3055689 | 10.22731667 | 1a | 10.23_301.2981n | 920 | 301.2981145 | 0.048094972 | in-house | M+H-H2O, M+H | C18H39NO2 | | - | - | - | - |
| 128 | 18:1(2S-OH) Ceramide | Sphingolipids | 540.5324906 | 22.67708333 | 2a | 22.68_539.5283n | 2098 | 539.5282848 | 1.040286764 | Mona | M+H-H2O, M+H | | | - | - | - | - |
| 129 | beta-Sitostenone | Steroid | 413.3777862 | 23.07665 | 2a | 23.08_412.3682n | 2179 | 412.3682117 | -5.588337899 | Metlin | M+H-H2O, M+H | C29H48O | | - | - | - | - |
| 130 | Similar to Campesterol | Sterol | 383.3673989 | 18.36516667 | 2a | 18.37_383.3674m/z | 1299 | | 0.427198389 | Metlin | M+H-H2O | C28H48O | | - | - | - | - |
| 131 | Similar to beta-Sitosterol | Sterol | 397.3828023 | 18.92661667 | 2a | 18.93_397.3828m/z | 1342 | | -0.182688015 | Metlin | M+H-H2O | C29H50O | | - | - | - | - |
| 132 | Similar to beta-Sitosterol | Sterol | 397.3829926 | 22.68193333 | 2a | 22.68_397.3830m/z | 2096 | | 0.276656079 | Metlin | M+H-H2O | C29H50O | | - | - | - | - |
| 133 | Similar to beta-Sitosterol | Sterol | 397.3819894 | 24.5802 | 2a | 24.58_397.3820m/z | 2378 | | -2.144401224 | Metlin | M+H-H2O | C29H50O | | - | - | - | - |
| 134 | Similar to Campesterol | Sterol | 383.3678492 | 22.22093333 | 1a | 22.22_383.3678m/z | 1927 | | 1.55474006 | in-house | M+H-H2O | C28H48O | | - | - | - | - |
| 135 | Similar to Stigmasterol | Sterol | 395.3676149 | 22.48326667 | 1a | 22.48_395.3676m/z | 154 | | 0.941260974 | in-house | M+H-H2O | C29H48O | | - | - | - | - |
| 136 | Similar to Stigmasterol | Sterol | 395.367335 | 18.66141667 | 2a | 18.66_395.3673m/z | 1321 | | 0.262575675 | in-house | M+H-H2O | C29H48O | | - | - | - | - |
| 137 | Stigmasterol | Sterol | 395.3672028 | 21.97138333 | 1a | 21.97_395.3672m/z | 1859 | | -0.060931309 | in-house | M+H-H2O | C29H48O | | - | - | - | - |
| 138 | Episterol | Sterol | 381.3512945 | 21.35953333 | 2a | 21.36_381.3513m/z | 1706 | | -0.708641603 | in-house | M+H-H2O | C28H46O | | - | - | - | - |
| 139 | Ergosterol | Sterol | 379.3359211 | 21.30553333 | 1a | 21.31_379.3359m/z | 1697 | | -0.011620532 | in-house | M+H-H2O | C28H44O | | - | - | - | - |
| 140 | STIGMASTA-4,22-DIEN-3-ONE-LIKE | | 411.3626232 | 22.89185 | 2a | 22.89_411.3626m/z | 2144 | | 1.171432081 | Metlin | M+H | C29H46O | | - | - | - | - |
| 141 | 4-acetoxy-(2h)-1,4-benzoxazin-3(4h)-one | | 178.050406 | 5.839533333 | 2a | 5.84_178.0504m/z | 37 | | 3.030441787 | Mona | M+H | C9H7NO3 | | - | - | - | - |
| 142 | 5S-HETE di-endoperoxide | | 425.2148277 | 16.21018333 | 2b | 16.21_425.2148m/z | 1164 | | 0.594100867 | Lipidmaps | M+Na | C20H34O8 | LMFA03000011 | - | - | - | - |
| 143 | 6-hydroxy-4a-(hydroxymethyl)-5-methyl-3-(1-methylethenyl)-3_4_5_6_7_8-hexahydronaphthalen-2-one | | 233.1539556 | 13.9182 | 2a | 13.92_233.1540m/z | 1045 | | 0.974782241 | in-house | M+H-H2O | C15H22O3 | | - | - | - | - |
